# TRPC3 is essential for functional heterogeneity of cerebellar Purkinje cells

**DOI:** 10.1101/478446

**Authors:** Bin Wu, Francois G.C. Blot, Aaron B. Wong, Catarina Osório, Youri Adolfs, R. Jeroen Pasterkamp, Jana Hartmann, Esther B. E. Becker, Henk-Jan Boele, Chris I. De Zeeuw, Martijn Schonewille

**Author notes:** Correspondence: Martijn Schonewille Postal address: Wytemaweg 80, 3015 CN, Rotterdam, the Netherlands.

## Abstract

Despite the canonical homogenous character of its organization, the cerebellum plays differential computational roles in distinct types of sensorimotor behaviors. However, the molecular and cell physiological underpinnings are unclear. Here we determined the contribution of transient receptor potential cation channel type C3 (TRPC3) to signal processing in different cerebellar modules. Using gain-of-function and loss-of-function mouse models, we found that TRPC3 controls the simple spike activity of zebrin-negative (Z–), but not of zebrin-positive (Z+), Purkinje cells. Moreover, *in vivo* TRPC3 also regulated complex spike firing and its interaction with simple spikes exclusively in Z– Purkinje cells. Finally, we found that eyeblink conditioning, related to Z– modules, but not compensatory eye movement adaptation, linked to Z+ modules, was affected in TRPC3 loss-of-function mice. Together, our results indicate that TRPC3 is essential for the cellular heterogeneity that introduces distinct physiological properties in an otherwise homogeneous population of Purkinje cells, conjuring functional heterogeneity in cerebellar sensorimotor integration.

## Introduction

Maintaining correct sensorimotor integration relies on rapid modifications of activity. The cerebellum is instrumental herein, evidenced by the fact that disruptions of cerebellar functioning, e.g. through stroke or neurodegenerative disorders, affect coordination and adaption of many types of behaviors such as gait, eye movements and speech^1,2^. The palette of behavioral parameters controlled by the cerebellum is also broad and includes features like timing^3-5^, strength^6,7^ as well as coordination^8,9^ of muscle activity. However, the pluriformity of behavioral features does not match with the homogeneity of the structure and cyto-architecture of the cerebellar cortex.

Recently, it has been uncovered that the sole output neurons of the cerebellar cortex, the Purkinje cells (PCs), can be divided into two main groups with a distinct firing behavior^10,11^. One group, consisting of PCs that are positive for the glycolytic enzyme aldolase C, also referred to as zebrin II^12,13^, shows relatively low simple spike firing rates, whereas the PCs in the other group that form zebrin-negative zones, fire at higher rates^10^. Zebrin II demarcates olivocerebellar modules, anatomically defined operational units each consisting of a closed loop between the inferior olive, parasagittal bands of the cerebellar cortex and the cerebellar nuclei^14,15^. Given that different motor domains are controlled by specific olivocerebellar modules^14,16^, the differential intrinsic firing frequencies may be tuned to the specific neuronal demands downstream of the cerebellum^17^. Thus, dependent on the specific behavior controlled by the module involved, the PCs engaged may show low or high intrinsic firing as well as related plasticity rules to adjust these behaviors.

Cellular heterogeneity can drive differentiation in the activity and plasticity of individual cells that operate within a larger ensemble^18^. The molecular and cellular determinants of differential electrophysiological processing in the cerebellar PC modules are just starting to be identified^19,20^. For example, while the impact of zebrin II itself is still unclear^10^, excitatory amino acid transporter 4 (EAAT4) and GLAST/EAAT1 can directly modulate simple spike activity of PCs as well as plasticity of their parallel fiber inputs in a zone specific manner^21,22^. Likewise, phospholipase C subtype β4 (PLCβ4) is required for climbing fiber elimination and PF-PC LTD through mGluR1 activation by spill-over glutamate and is only expressed in zebrin-negative modules^23-25^. The alpha isoform of mGluR1 (mGluR1a) is responsible for PLCβ4 activation and is uniformly expressed in cerebellar PCs^26^. Conversely, the mGluR1b receptor is expressed in a pattern complementary to that of zebrin^27^, but it is less clear to what extent mGluR1b may affect PCs.

Given that mGluR1b interacts with TRPC3 to drive mGluR1-dependent currents^28^, we set out to test the hypothesis that TRPC3 is a key player in the molecular machinery responsible for differential control over PC activity and function. We demonstrate that TRPC3 in the brain has particularly strong expression in the cerebellum, in a pattern complementary to zebrin in the vermis and more uniform in the hemispheres. We examined the impact of TRPC3 gain-of-function and loss-of-function mutations and found effects on the spiking rate of Z– but not Z+ PCs *in vitro*. *In vivo* recordings during quiet wakefulness in the same mutants revealed that the level of TRPC3 influences both simple spike and complex spike rates, and the interaction between the two, also selectively in Z– modules. Finally, we show that adaptation of compensatory eye movements, which is controlled by Z+-modules in the vestibulocerebellum^10,29^, is not affected by the loss of TRPC3 function, whereas the learning rate during eyeblink conditioning, which is linked to the Z– modules^30,31^, is decreased after PC-specific ablation of TRPC3, highlighting the behavioral relevance of firing rate modulation by TRPC3.

## Results

### Specific expression pattern and subcellular localization of TRPC3 in the mouse brain

The expression of TRPC3 and how it relates to that of zebrin in the adult mammalian brain is unclear, in part due to poor antibody quality. Using a novel TRPC3-specific antibody (Cell signaling, #77934), we examined the immunohistochemistry of TRPC3. We found that in the mouse brain TRPC3 is most prominently expressed in the cerebellum (**Fig. 1a**), specifically in PCs and unipolar brush cells (UBCs) (**Fig. 1b**). However, upon further scrutiny, it is clear that, although expressed in all PCs, endogenous TRPC3 does not distribute homogeneously. TRPC3 expresses in a pattern that in the vermis complements that of zebrin, while in the hemispheres it appears more uniform (**Fig. 1b-c, Supplementary Fig. 1, 2**). Specifically, the TRPC3 level, in the anterior cerebellum (referred to as lobules I-III), where the PCs are predominantly Z–, is quite intense; while less so in the posterior PCs (referred to as lobule X), which are primarily Z+ (**Supplementary Fig. 2a-b**). Although clearly observable in our standard immunohistochemistry, this pattern was visualized in a more comprehensive manner using whole-mount brain light sheet imaging. The antibody staining appears to be of better quality in the iDISCO protocol. The lens and resolution of the light sheet also circumvent stitching artefacts resulting from a tile scanning of the confocal microscope by imaging the full cerebellum in one single scanning window (**Supplementary Movie**). The anterior and posterior differences of the protein amount were confirmed by western blot analysis (**Supplementary Fig. 3a-b**).

**Fig.1.**
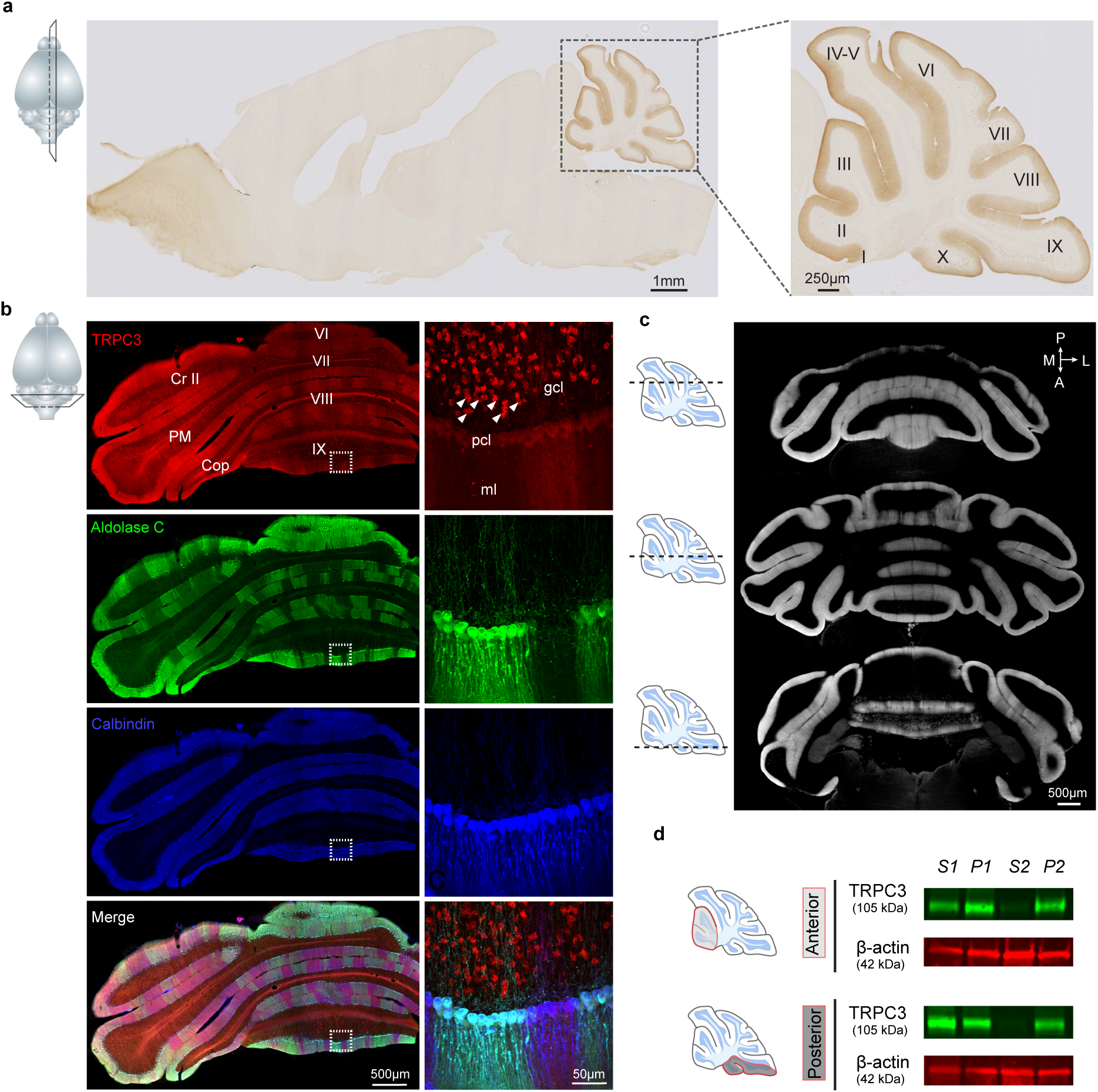
TRPC3 is predominantly expressed in the cerebellum in a zebrin-related pattern. **a**, Representative image and magnification (right) of sagittal cryosection of an adult mouse brain stained with anti-TRPC3. Inset, plane of section. **b**, Coronal immunofluorescence images with anti-TRPC3 (red), anti-aldolase C (green) and anti-calbindin (blue) staining of the cerebellar cortex (left), with magnifications (right). TRPC3 is expressed in the cerebellar PCs and UBCs (triangles), in a pattern that in the vermis complements that of zebrin and appears more uniform in the hemispheres. Inset, plane of section. **c**, Individual images of a light sheet imaging-based reconstruction of a mouse brain cleared with iDISCO and stained with anti-TRPC3. Three different planes (insets) of whole cerebellum show sagittal compartmentations across lobules. **d**, Immunoblots of TRPC3 by using synaptic protein extraction protocol on the anterior (top) and posterior (bottom) cerebellum. TRPC3 is present in the homogenate (S1) and enriched in the membrane (P1) and synaptosomes (P2), but not in the cytosol (S2). I-X, cerebellar lobules I-X; Cr II, Crus II; PM, paramedian lobule; Cop, Copula Pyramidis; gcl, granule cell layer; pcl, Purkinje cell layer; ml, molecular layer; D, dorsal; V, ventral; M, medial; L, lateral.

**Fig.2.**
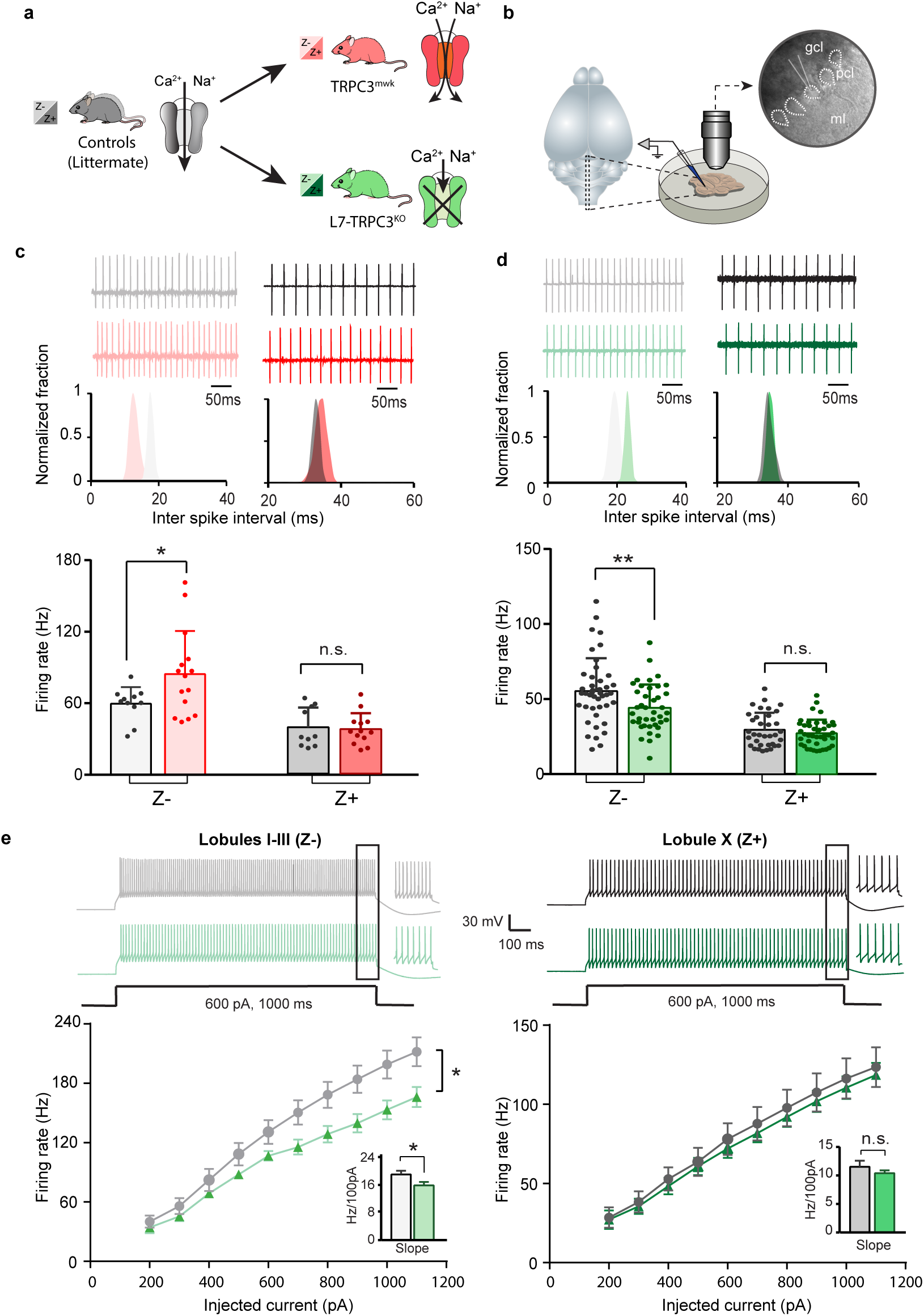
Differential controls of PCs firing properties by TRPC3 *in vitro.* **a**, Schematic drawing of TRPC3 channel function in control (black), gain-of-function (TRPC3^Mwk^, red) and loss-of-function (L7-TRPC3^KO^, green) mice. **b**, Schematic approach illustrating of PCs (right circle, dashed lines) recording *in vitro*, in acute sagittal slices (left). **c**,**d**, Representative traces of cell-attached PC recordings (top) and corresponding inter spike interval (ISI) distributions (middle) in a Z– PC (left) and a Z+ PC (right) of TRPC3^Mwk^ (**c**) and L7-TRPC3^KO^ (**d**) mice. Z– PCs were affected in TRPC3^Mwk^ (**c**, light-red, n=15 cells/N=4 mutant mice vs. n=11 cells/N=2 littermate controls, *t*_19_=-2.43, *P*=0.025 and in L7-TRPC3^KO^ mice (**d**, light-green, n=40/N=6 mutants vs. n=43/N=5 controls, *t*_81_=2.69, *P*=0.009). No differences in the firing rate of Z+ PCs in TRPC3^Mwk^ (**c**, dark-red, n=13/N=4 mutants vs. n=10/N=2 controls, *t*_21_=0.242, *P*=0.811) and L7-TRPC3^KO^ mice (**d**, dark-green, n=36/N=10 mutants vs. n=35/N=4 controls, *t*_64_=0.937, *P*=0.352). **e**, Whole-cell patch-clamp recordings in slice from PCs of L7-TRPC3^KO^ and control mice were used to test intrinsic excitability, by keeping cells at a holding potential of −65 mV and evoking action potentials by current steps of 100 pA (example, top). Top, exemplary traces evoked by current injection at 600 pA. Bottom, Input-output curves from whole-cell recordings of L7-TRPC3^KO^ mice of Z– PCs (left, n=17/N=5 mutants vs n=17/N=5 controls, t_32_=-2.20, P=0.035) and Z+ PCs (right, n=12/N=5 mutants vs n=12/N=4 controls, t_22_=-0.95, P=0.354). gcl, granule cell layer; pcl, Purkinje cell layer; ml, molecular layer. **c-d**, data are represented as mean ± s.d.; **e**, data are represented as mean ± s.e.m.. For values see **Supplementary Table 2**.

**Fig.3.**
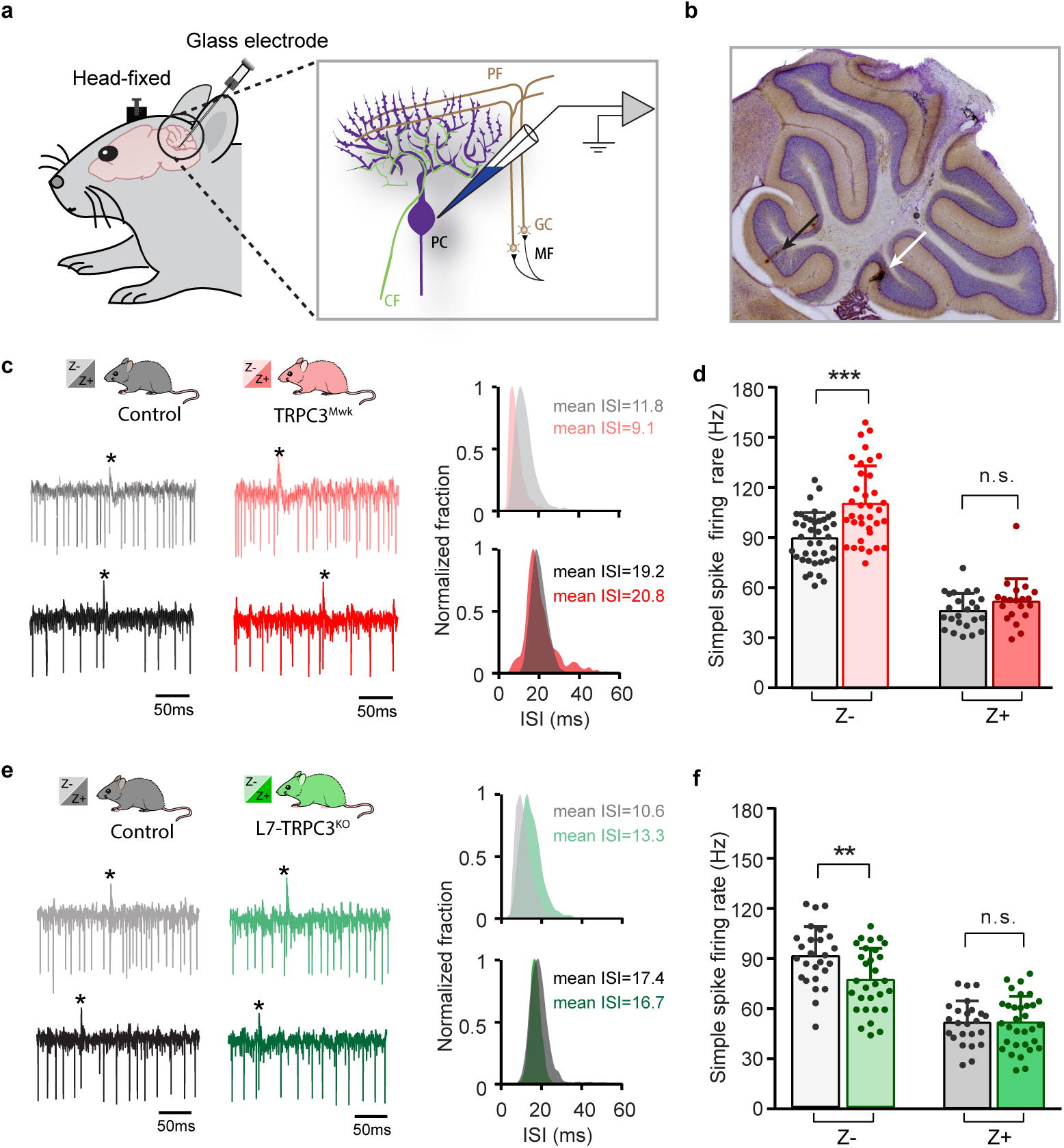
*In vivo* simple spike firing rate of Z–, but not Z+ PCs is controlled by TRPC3. **a**, Schematic illustration of extracellular recording configuration *in vivo*. PF, parallel fiber; CF, climbing fiber; MF, mossy fiber; GC, granule cell. **b**, Representative sagittal cerebellar section with recording sites labelled by BDA injection, in lobule II (black arrow) and X (white arrow). **c**, Representative example traces (left) and ISI distributions (right) of a Z– PC (top) and a Z+ PC (bottom) in gain-of-function TRPC3^Mwk^ mice. **d**, PC simple spike firing rate recorded *in vivo* in TRPC3^Mwk^ mice compared to control littermates, for the Z– lobules I-III (light-red, n=36/N=7 mutants vs. n=40/N=6 controls, *t*_60_=-4.58, *P*<0.001) and the Z+ lobule X (dark-red, n=20/N=6 mutants vs. n=24/N=5 controls, *t*_42_=-1.47, *P*=0.148). **e**, Representative example traces (left) and ISI distributions (right) in a Z– PC (top) and a Z+ PC (bottom) of loss-of-function L7-TRPC3^KO^ mice. **f**, PC simple spike firing rate of L7-TRPC3^KO^ mice compared to controls, for Z– lobules I-III (light-green, n=30/N=7 mutants vs. n=26/N=8 controls, *t*_54_=2.88, *P*=0.006) and in Z+ lobule X (dark-green, n=32/N=8 mutants vs. n=24/N=6 controls, *t*_54_=-0.053, P=0.958). Data are represented as mean ± s.d., for values see **Supplementary Table 3**.

Our immunohistochemical imaging reveals that TRPC3 is present in the soma and dendritic arbor of PCs (**Fig. 1b, Supplementary Fig. 2a-b**), to further examine the subcellular localization of TRPC3 in the cerebellum, we performed immunoblots of isolated fractions following a synaptic protein extraction procedure (**Supplementary Fig. 3c**). As expected, TRPC3, a channel protein, is abundantly present in the membrane and almost completely absent in the cytosol (**Supplementary Fig. 3d**). Moreover, TRPC3 is enriched in synapstosomes (**Supplementary Fig. 3d**), in line with the common conception of mGlur1b-dependent activation^26,28^. Together, these results indicate that, within the brain, strong TRPC3 expression is restricted to the cerebellum, where it is present in all PCs and UBCs, but at particularly high levels in Z– PCs.

### TRPC3 differentially controls the physiological properties of PCs *in vitro*

Next, we investigated the contribution of TRPC3 to cerebellar function in Z+ and Z– PCs using both loss-of-function and gain-of-function mouse models (**Fig. 2a**). TRPC3-Moonwalker (TRPC3^Mwk^) mice harbor a point mutation resulting in TRPC3 gain-of-function through increased Ca^2+^ influx upon activation^32^. Inversely, TRPC3 was selectively ablated from cerebellar PCs by crossing mice carrying *loxP*-flanked TRPC3 alleles^28^ with L7-Cre (*PCP2*-Cre)^33^ mice, generating L7-TRPC3^KO^ mice. Western blotting and immunostaining of the anterior (Z–) and the posterior (Z+) cerebellar cortex of L7-TRPC3^KO^ mice confirmed that protein levels are reduced (**Supplementary Fig. 3, 4**). The loss of TRPC3 was specific for cerebellar PCs, as TRPC3 expression in UBCs was not affected (**Supplementary Fig. 4b,** white arrow heads).

**Fig.4.**
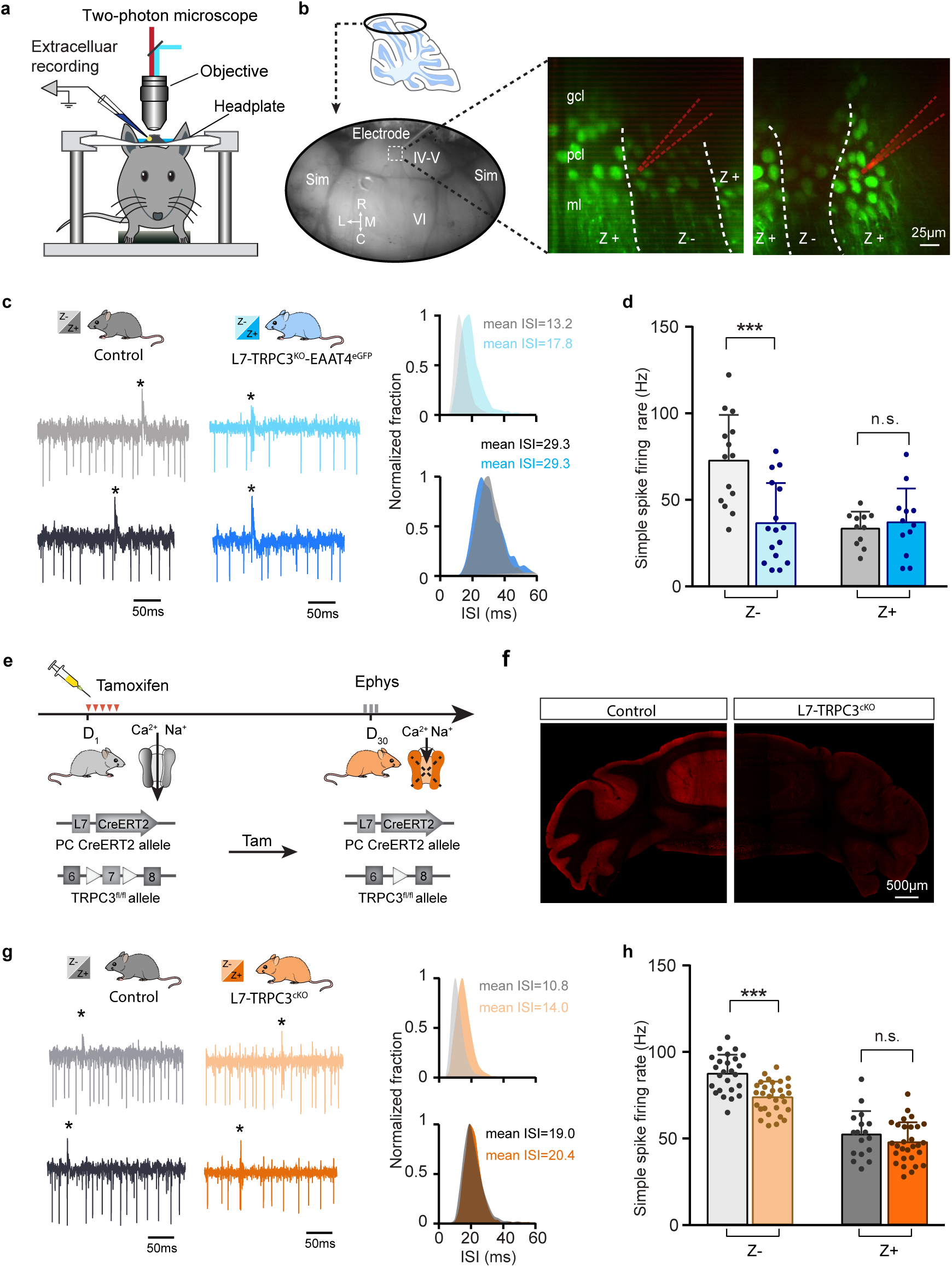
TRPC3 effects follow zebrin-identity and are not developmental. **a**, Schematic experimental setup for two-photon imaging-based targeted PC recordings, *in vivo*. **b**, Sagittal view of cerebellum (schematic, top) indicating the recording region in the ellipse (bottom). Representative images (right) show the visualization of Z+ bands (strong green) in an awake L7-TRPC3^KO^-EAAT4^eGFP^ mouse, with recording electrodes (red) positioned in Z– (left) and Z+ (right) bands. **c**, Representative firing traces (left) and ISI distributions (right) in a Z– PC (top) and a Z+ PC (bottom) of loss-of-function L7-TRPC3^KO^-EAAT4^eGFP^ mice (blue) and control littermates (no Cre; gray). **d**, Average simple spike firing rate of PCs recorded from adjacent modules of L7-TRPC3^KO^-EAAT4^eGFP^ mice and those in control littermates. Comparison for Z– PCs (light-blue, n=16/N=3 mutants vs. n=14/N=2 controls, *t*_28_=3.99, *P*<0.001), and Z+ PCs (dark-blue, n=12/N=3 mutants vs. n=12/N=2 controls, *t*_21_=-0.550, *P*=0.588). **e-f**, Intraperitoneal tamoxifen injections for five days (D_1-5_) to trigger TRPC3 gene ablation solely in PCs (using L7 promotor) in adult L7^CreERT2^-TRPC3^fl/fl^ mice. Open triangles indicate *loxP* sites. PC in vivo extracellular activity was recorded four weeks later (D_29-31_) in L7-TRPC3^cKO^ mice (orange). TRPC3 deletion was confirmed after experiment by confocal image using anti-TRPC3 staining (**f**). **g**, Representative firing traces (left) and ISI distributions (right) in a Z– PC (top) and a Z+ PC (bottom) of L7-TRPC3^cKO^ mice. **h**, Simple spike firing rate *in vivo* in L7-TRPC3^cKO^ and control mice (no Cre). Comparison for Z– PCs (light-orange, n=30/N=4 mutants vs. n=25/N=4 controls, *t*_53_=5.05, *P*<0.001), and Z+ PCs (dark-orange, n=29/N=4 mutants vs. n=17/N=3 controls, *t*_44_=1.21, *P*=0.234). Sim, simplex lobule; IV-VI, lobules IV-VI, R, rostral, C, caudal; L, lateral, M, medial. Data are represented as mean ± s.d., for values see **Supplementary Table 3**.

PCs are intrinsically active pace-making neurons, which fire regular action potentials even when deprived of synaptic inputs^34,35^. To determine the contribution of TRPC3 to the activity of Z+ and Z– PCs, we performed *in vitro* electrophysiological recordings on sagittal sections of adult mice of both mutants (**Fig. 2b**), taking lobules X and I-III as proxies for Z+ and Z– PC modules, respectively (see ref.^10,15^). In littermate controls, the intrinsic firing rate of Z– PCs is higher than that of Z+ PCs, confirming previous results^10^. Gain-of-function TRPC3^Mwk^ mice showed a decrease in inter spike intervals (ISI) and an increase in PC simple spike firing rate selectively in Z– PCs, without affecting Z+ PCs (**Fig. 2c**). Inversely, ablating TRPC3 from PCs caused an increase in ISI and decrease in firing rate in Z– PCs, again without affecting Z+ PCs (**Fig. 2d**). We also assessed the regularity of firing activities by measuring the coefficient of variation (CV) and the coefficient of variation of adjacent intervals (CV2) of ISI. Both the CV and CV2 of Z– PCs in lobules I-III declined significantly in L7-TRPC3^KO^ mice, while remaining unchanged in TRPC3^Mwk^ mice; in contrast, in Z+ lobule X, none of these parameters were altered in either TRPC3^Mwk^ or L7-TRPC3^KO^ mice (**Supplementary Fig. 5**).

**Fig.5.**
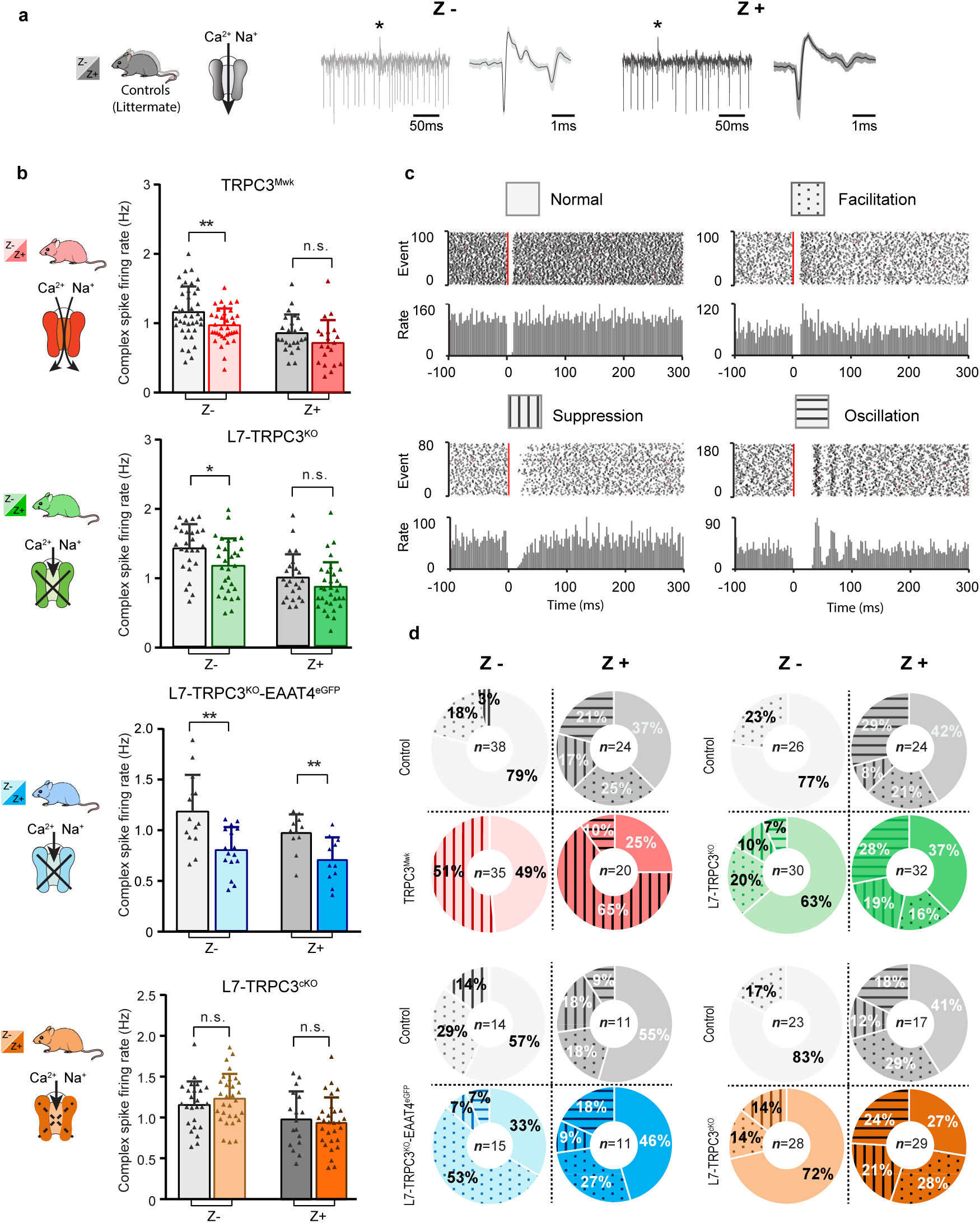
Complex spikes and complex spike - simple spike interaction are affected by TRPC3 mutations. **a**, Representative PC recording traces and complex spikes shape of Z– (light black) and Z+ (dark black) PCs in the control mice. **b**, Top half, comparison of complex spike firing rates in TRPC3^Mwk^ (red) and L7-TRPC3^KO^ (green) mice versus their respective littermate controls for Z– PCs (TRPC3^Mwk^: *t*_68_=2.68, *P*=0.009; L7-TRPC3^KO^: *t*_54_=2.50, *P*=0.016) and Z+ PCs (TRPC3^Mwk^: *t*_42_=1.56, *P*=0.126; L7-TRPC3^KO^: *t*_54_=1.41, *P*=0.164). Bottom half, comparison of complex spike firing rates in L7-TRPC3^KO^-EAAT4^eGFP^ (blue) and L7-TRPC3^cKO^ (orange) mice versus their respective controls for Z– PCs (L7-TRPC3^KO^-EAAT4^eGFP^: *t*_28_=3.49, *P*=0.002; L7-TRPC3^cKO^: *t*_53_=-0.940, *P*=0.352) and Z+ PCs (L7-TRPC3^KO^-EAAT4^eGFP^: *t*_20_=3.03, *P*=0.007; L7-TRPC3^cKO^: *t*_44_=0.448, *P*=0.656). **c**, Raster plots of simple spike activity around the occurrence of each complex spike (−100 to +300 ms). These peri-complex splike time histograms can, based on post-complex spike activity, be divided into one of four types: normal (no change), facilitation, suppression and oscillation. **d**, The distribution of post-complex spike response types for Z– and Z+ PCs, in TRPC3^Mwk^, L7-TRPC3^KO^, L7-TRPC3^KO^-EAAT4^eGFP^ and L7-TRPC3^cKO^ mice. Data are represented as mean ± s.d., for values see **Supplementary Table 3**.

To verify the effect of TRPC3 deletion on other cell physiological properties of PCs, we performed whole-cell patch-clamp recordings in a subset of PCs. Injections of current steps into PCs evoked increasing numbers of action potential, in the presence of blockers for both excitatory and inhibitory synaptic inputs. In line with the cell-attached recordings, in loss-of-function L7-TRPC3^KO^ mice, PC intrinsic excitability, quantified by the slope of firing rate versus current injection curve, was significantly reduced in lobules I-III, but unchanged in lobule X, compared with those of littermate controls (**Fig. 2e**). Other physiological parameters in terms of holding current, amplitudes, half-widths and after-hyperpolarization amplitudes, were not significantly affected in either lobules I-III or lobule X (**Supplementary Fig. 6**).

**Fig.6.**
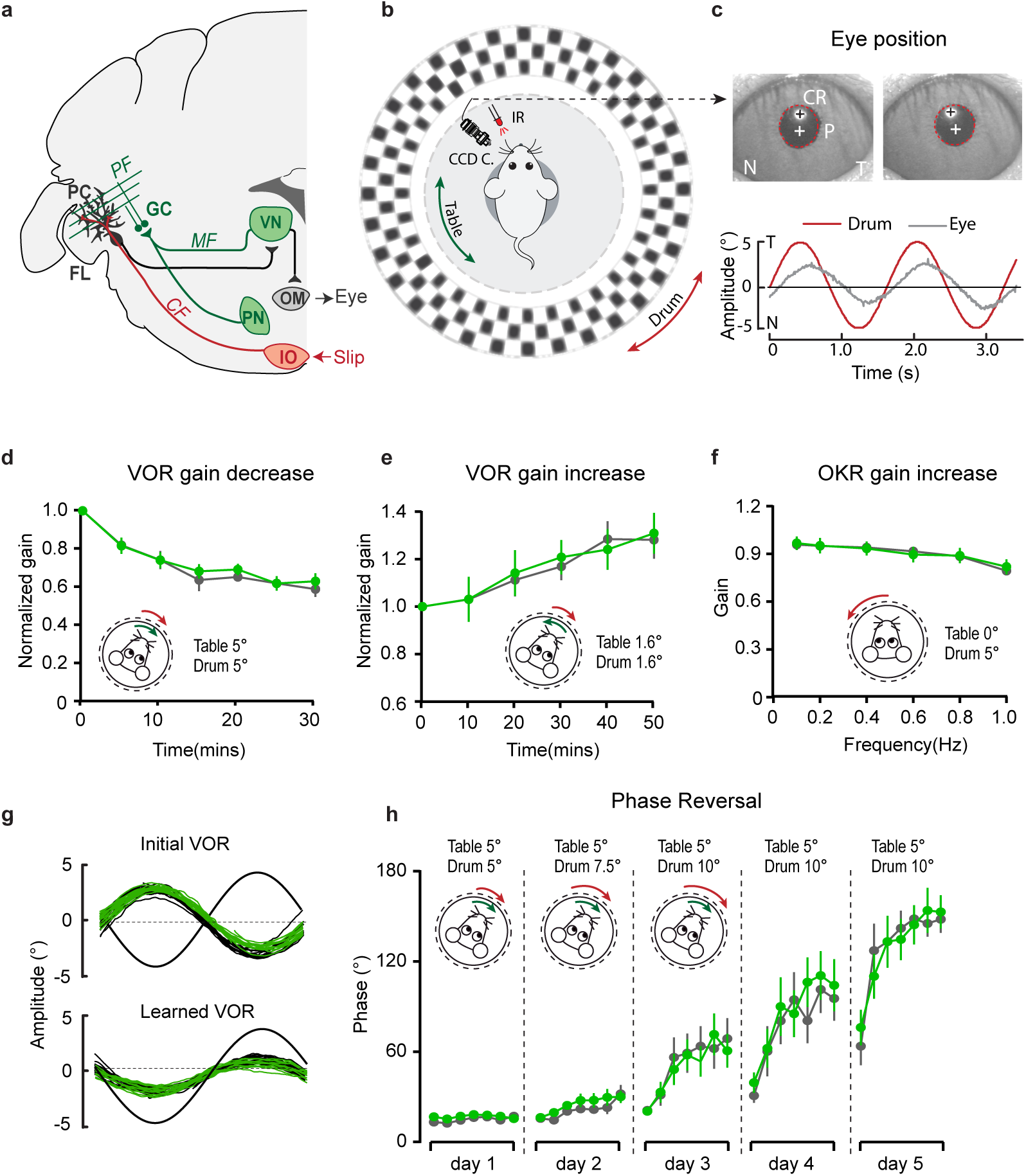
PC-specific deletion of TRPC3 does not affect Z+-dependent VOR adaptation. **a**, Cerebellar circuitry controlling compensatory eye movements and their adaptation. PCs in the flocculus (FL) receive vestibular and visual input via the mossy fiber (MF) - parallel fiber (PF) system (green) and climbing fiber input (CF, red) from the inferior olive (IO), indicating retinal slip. These two inputs converge on PCs, which influence eye movements via the vestibular nuclei (VN) and the oculomotor (OM) neurons. PN, pontine nuclei; GC, granule cell. **b**, Schematic illustration of eye movement recording setup. Mice are head-fixed in the center of a turntable for vestibular stimulation and surrounded by a random dotted pattern (‘drum’) for visual stimulation. A CCD camera was used for infrared (IR) video-tracking of the left eye. **c**, Top, examples of nasal (N) and temporal (T) eye positions. Red circles, pupil fit; black cross, corneal reflection (CR); white cross, pupil center. Bottom, example trace of eye position (grey) with drum position (red), during stimulation at an amplitude of 5° and frequency of 0.6 Hz. **d**, L7-TRPC3^KO^ and control mice were subjected to six 5-min training sessions with mismatched in-phase visual and vestibular stimulation (in light, see insets), aimed at decreasing the VOR gain (probed in the dark before, between and after sessions). **e**, Similar, but now mice were trained with out-of-phase stimulation, aimed at increasing VOR gain. **f**, Re-recording of OKR gain following the VOR phase reversal training (see **g-h**) to test OKR gain increase (compare to **Supplementary Fig. 12, left**). **g**, Multiple-day training using in-phase mismatch stimulation (see inset in **h**) aimed at reversing the direction of the VOR (quantified as a reversal of the phase). Representative eye position recordings of VOR before (top) and after (bottom) training. **h**, Results of five days of VOR phase reversal training, probed by recording VOR (in the dark before, between and after sessions) with mice kept in the dark in overnight. Data are represented as mean ± s.e.m., N=11 mutants versus N=13 controls, all P > 0.05, ANOVA for repeated measurements. See **Supplementary Table 4** for values.

Together, our *in vitro* recordings from gain- and loss-of-function mutants indicate that TRPC3 selectively controls the activity in Z– PCs, without affecting excitability or other cell intrinsic properties. Thus, at least *in vitro*, TRPC3 contributes to the difference in intrinsic firing activity between Z+ and Z– PCs, by directly controlling the intrinsic excitability of Z– PCs.

### TRPC3 regulates the activity of simple spikes selectively in Z– PCs *in vivo*

To examine the role of TRPC3 in the closed loop, intact cerebellar module, we next performed PC recordings *in vivo* in adult mice during quiet wakefulness (**Fig. 3a**). PCs could be identified during extracellular recordings by the presence of complex spikes, while the consistent presence of a pause in simple spikes following each complex spike confirmed that the recording was obtained from a single unit^36^. PC recording locations in either Z– lobules I-III or Z+ lobule X were confirmed with iontophoretic injections of biotinylated dextran amine (BDA), which could be identified by immunostaining (**Fig. 3b**). As we showed before^10,37^, PCs in Z– modules fired simple spikes at a higher rate than those in Z+ modules (**Fig. 3d,f**).

*In vivo*, in the presence of physiological inputs the PCs in Z– lobules I-III of TRPC3^Mwk^ mutants showed a decreased ISI and increased simple spike firing rate, whereas the Z+ PCs were unaffected. Conversely, Z– PCs in L7-TRPC3^KO^ mutants featured an increased ISI and decreased simple spike firing rate, but again without changes in PCs of the Z+ lobule X, all compared to those of their littermate controls (**Fig. 3c-f**). Unlike *in vitro*, PCs in the L7-TRPC3^KO^ mice showed CV and CV2 comparable to controls for both Z– and Z+ modules (**Supplementary Fig. 7g-i**). The CV of simple spike ISI was, however, prominently elevated in both Z– and Z+ modules in TRPC3^Mwk^ mutants (**Supplementary Fig. 7a-c**). It should be noted that PC regularity *in vivo* is largely determined by external inputs (compared **Supplementary Fig. 5** to **7**), which thereby can offset those intrinsic variations induced by the mutation of TRPC3. The irregular firing activity of PCs in TRPC3^Mwk^ mutants, at least for Z+ PCs, may be attributed to impaired function or degeneration of UBCs, while the physiological synaptic input *in vivo* in L7-TRPC3^KO^ mice could obscure the regularity changes observed *in vitro* in these mice.

**Fig.7.**
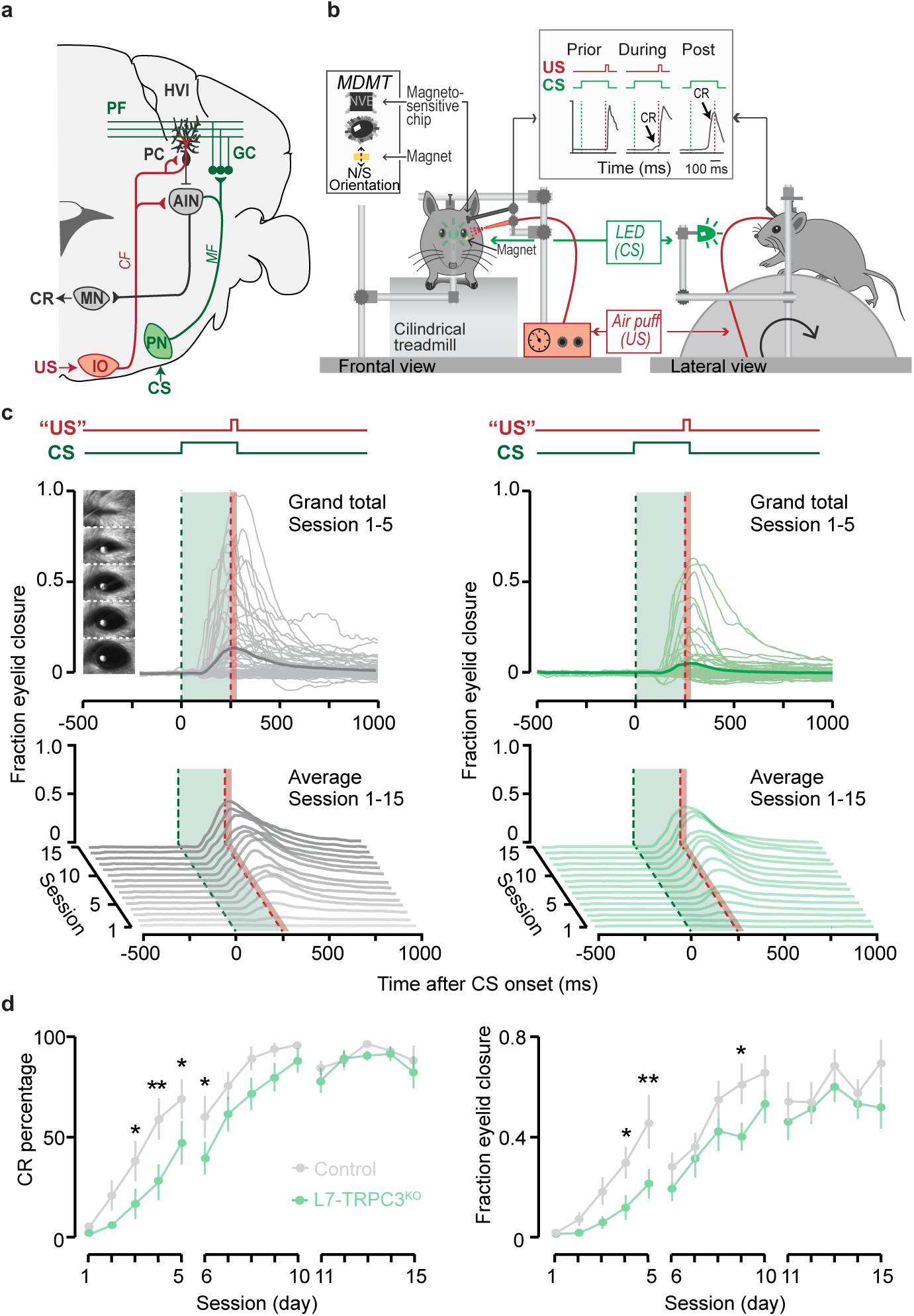
Eyeblink conditioning, linked to Z– modules is delayed in L7-TRPC3^KO^ mice. **a**, Cerebellar circuitry controlling eyeblink conditioning. PCs in the paravermal region around the primary fissure receive inputs carrying sensory information from e.g. the pontine nucleus (PN) through the MF-PF pathway and the error signal from the inferior olive (IO) through the climbing fibers (CF). These PCs in turn influence eyelid muscles via the anterior interposed nucleus (AIN) and motor nuclei (MN). **b**, Schematic illustration of eyeblink conditioning setup. Head fixed mice on a freely moving treadmill, are presented a green LED light (conditional stimulus, CS) followed several hundred milliseconds later by a weak air-puff on the eye (unconditional stimulus, US). As a result of repeated CS-US pairings, mice will eventually learn to close their eye in response to the CS, which is called the conditioned response (CR). Eyelid movements were recorded with the magnetic distance measurement technique (MDMT). **c**, Comparison of fraction of eyelid closure between controls (left) and L7-TRPC3^KO^ mice (right). Top, session averages (thin-lines) per mouse and overall average (thick-lines) for the first 5 days. Insets: mouse eye video captures show eyelid closure ranging from 0 (fully-open) to 1 (fully-closed). Bottom, waterfall plot of the averaged eyeblink trace during CS-only trials for the 15 daily sessions. **d**, The CR percentage and CR amplitude for L7-TRPC3^KO^ mice initially have an significantly slower acquisition but eventually reach the same levels as control littermates. Data are represented as mean ± s.e.m., N=15 mutants versus N=15 controls, *P* values were all FDR corrected for multiple comparisons, see **Supplementary Table 5** for values and statistics.

In short, even *in vivo*, in the presence of all physiological inputs both gain-of-function and loss-of-function mutations of TRPC3 exclusively affects Z– PCs, with the most pronounced, persistent effect being the mutation-selective influence on simple spike firing rate.

### TRPC3-related effects correlate with zebrin expression and are independent of development

Our results so far have identified selective TRPC3-related effects by comparing lobules I-III and X, as proxies for Z– and Z+ modules. Immunohistochemical analysis indicated that the TRPC3 expression differs substantially between these lobules (**Supplementary Fig. 1, Supplementary Movie**), suggesting that the effects of gain- and loss-of-function mutations could be directly related to protein levels. Alternatively, other differences in molecular machinery could underlie or further enhance this cellular differentiation, for instance through mGluR1b-related effects. As the difference in TRPC3 expression is minimal or absent in the more lateral parts of the cerebellum (**Supplementary Fig. 1, 2)**, recording the activity of adjacent Z– and Z+ PCs there would solve this question. To this end, we crossed L7-TRPC3^KO^ mice with EAAT4^eGFP^ mice that express eGFP in Z+ PCs to generate L7-TRPC3^KO^-EAAT4^eGFP^ mice (**Supplementary Fig. 8**). Using two-photon microscopy, we identified Z+ and Z– modules on the dorsal surface (lobules IV-VI and simplex) of the cerebellum and recorded PC activity (**Fig. 4a-b**). Here, the absence of TRPC3 attenuated the firing rate and enhanced the irregularity of Z– PCs even more robustly, without an effect on Z+ PCs (**Fig. 4c-d, Supplementary Fig. 9a-c**, cf **Fig. 3f**). These results argue against a direct link between simple spike firing rate and TRPC3 levels and support the concept that other proteins, e.g. mGluR1b, influence TRPC3 activity and thereby control the spiking activity of PCs.

To test the possibility that developmental effects influenced PC activity in the adult mice we tested (L7^Cre^ begins to be expressed in postnatal week 1-2), we crossed the *loxP*-flanked TRPC3 mice with tamoxifen-dependent L7^Cre-ERT2^ to generate L7-TRPC3^cKO^ mice (**Fig. 4e**). If the divergent effects of TRPC3 on Z– and Z+ PCs are completely or in part of developmental origin, we should observe no or less changes in L7-TRPC3^cKO^ adult mice after tamoxifen injections (injected after maturation). Four weeks after tamoxifen treatment, L7-TRPC3^cKO^ mice showed a virtually complete ablation of TRPC3 in PCs (**Fig. 4f**). Again, simple spike regularity and firing rate were affected in Z–, but not Z+ PCs of tamoxifen injected adult L7-TRPC^cKO^ mice (**Fig. 4g-h, Supplementary Fig. 9g-i**), in a manner similar to that in L7-TRPC^cKO^.

Taken together and combined with L7-TRPC3^KO^ data, these results indicate that the TRPC3-dependent effects in zebrin-identified PCs are independent of cerebellar development or developmental compensation. Moreover, the larger effect of TRPC3 ablation on Z– PCs in areas where its expression is similar to that in Z+ PCs suggests that other proteins contribute to the state of increased excitability in Z– PCs.

### TRPC3 mutations selectively affect the activity in Z– olivocerebellar modules

PCs in the cerebellar cortex, form a closed loop with the cerebellar nuclei neurons they innervate by their axon output and the olivary neurons from which they receive their climbing fiber input^14^. If TRPC3 contributes to the output of this loop, one could hypothesize that other elements in the loop should be affected by the mutations^7,38^. To test this hypothesis, we examined complex spikes activity in PCs, as the complex spike directly reflects the activity of the climbing fiber and thereby that of the inferior olivary neuron it originates from^38^. We identified complex spikes based on their characteristic shape in our *in vivo* recordings from Z– lobules I-III or Z+ lobule X (**Fig. 5a**). Complex spike firing rates were, similar to simple spike rates, higher in Z– than in Z+ PCs (**Fig. 5b**), as shown previously^10^. Chronic manipulations of TRPC3 activity, gain- and loss-of-function, in PCs predominantly affected complex spike firing rate in Z–, but not Z+ PCs (**Fig. 5b**). Intriguingly, acute ablation of TRPC3 in L7-TRPC3^cKO^ mice did not affect complex spike activity in terms of firing rate, CV, CV2 or pause in simple spikes following climbing fiber activation (CF-pause) in Z– PCs (**Fig. 5b, Supplementary Fig. 9j-l**). In line with the lower simple spike firing rates in loss-of-function TRPC3 mutants, the CF-pause of L7-TRPC3^KO^ and L7-TRPC3^KO^-EAAT4^eGFP^ mice were longer, selectively in Z– PCs (**Supplementary Fig. 7l, S9f**). Except for the CV value, other complex spike parameter changes in TRPC3^Mwk^ mice were not affected (**Supplementary Fig. 7d-f**). Together, *in vivo* experiments indicate that TRPC3 also selectively affects the activity in the inferior olive in that the Z– modules are most prominently affected, and this influence has a developmental component.

Complex spikes are known to have a direct influence on simple spike activity (CS-SS)^10,39^. Based on the peri-complex spike time histograms, we could categorize four different types of simple spike responses following the CF-pause (see also ref.^10^), including no change in rate (normal), increased simple spike activity (facilitation), decreased simple spike activity (suppression), and a superimposed oscillatory pattern (oscillation) (**Fig. 5c**). Our data confirmed our previous finding^10^ that the CS-SS interaction pattern among the Z+ and Z– PCs is different in that the facilitation prevails in the Z− PCs, whereas the suppression and oscillation types occur predominantly in the Z+ PCs (**Fig. 5d**). In addition, we found that manipulation of TRPC3 activity changed the types of CS-SS responses most frequently in Z– PCs (**Fig. 5d**). Interestingly, Z– PCs exhibited much more suppression in gain-of-function TRPC3^Mwk^ mutants and vice versa more facilitation in loss-of-function L7-TRPC3^KO^-EAAT4^eGFP^ mice, compared to those in their littermate controls (**Fig. 5d**), suggesting that Z– PCs partly compensate for the effects of TRPC3 manipulation.

Together, these results indicate that TRPC3 controls not only the activity of PCs, but also that of the inferior olivary neurons, another element in the olivocerebellar loop. Moreover, manipulation of TRPC3 activity alters the interaction between complex spikes and simple spikes.

### Functional heterogeneity of TRPC3 is reflected in differential effects on motor behaviors

The ultimate question is: does cellular heterogeneity of PCs also differentially affect their contribution to specific cerebellar functions? As the TRPC3^Mwk^ mutation is not cell-specific and affects for instance also UBCs, we focused on the behavioral effects in L7-TPRC3^KO^ mice. Before testing specific functions, we first evaluated the consequences of the manipulations of TRPC3 on locomotion, a type of behavior that by nature requires the entire body and as such can be linked to many sub-regions of the cerebellar cortex, from the Z+ vestibular zones to the Z– anterior lobules. We first investigated whether these mutant mice showed any obvious deficits in locomotion using the Erasmus Ladder^9^. L7-TPRC3^KO^ mice could not be discriminated from control littermates by the percentage of different types of steps, including lower steps, also known as missteps (**Supplementary Fig. 10**). The apparent discrepancy with earlier evidence in stride width in the global TRPC3 knockout^28^ could be due to the different methods or the fact that UBCs, particularly important in the vestibular zone, are also affected in that mouse model^40^.

Next, we subjected L7-TPRC3^KO^ mice to two specific, but intrinsically distinct types of cerebellum-dependent learning tasks, i.e., vestibulo-ocular reflex (VOR) adaptation and eyeblink conditioning. VOR adaptation is the adjustment of the amplitude and/or direction of compensatory eye movements controlled by the vestibulocerebellum (**Fig. 6a-c**), which is predominantly Z+ (**Supplementary Fig. 11a**). Eyeblink conditioning requires the animal to generate a well-timed movement following a previously unrelated sensory input and is linked to more anterior regions that are largely Z– (**Fig. 7a, Supplementary Fig. 11b**). Note that the difference in zebrin labeling is pronounced between the two related regions, while the difference in TRPC3 staining is less clear (**Supplementary Fig. 11**). Nonetheless, given the electrophysiological changes described above, we hypothesized that altered TRPC3 function should impair Z– linked eyeblink conditioning, whereas VOR adaptation would be unaffected.

Before examining adaptation, we first tested if the basal eye movement reflexes, the optokinetic reflex driven by visual input (OKR) and the vestibular input-driven VOR (in the dark) and visually-enhanced VOR (VVOR, in the light), were affected. Neither the gain (the ratio of eye movement to stimulus amplitude), nor the phase (timing of the response relative to input), differed significantly between L7-TPRC3^KO^ mutants and littermate controls (**Supplementary Fig. 12a**). Next, using mismatched visual and vestibular stimulation, we tested the ability of mutant mice to adapt their compensatory eye movements. When L7-TPRC3^KO^ mice were subjected to both out-of-phase and in-phase training paradigms, we did not observe any significant deficit in the VOR gain increase and VOR gain decrease, respectively (**Fig. 6d-e**). To evaluate the ability of the mice to perform a long-term, more demanding adaptation, we subjected the mice for three more days, following the gain decrease training, to a training stimulus aimed at reversing the direction of their VOR, referred to as VOR phase reversal (**Fig. 6g**). Again, no difference was found between L7-TPRC3^KO^ and control littermate mice: neither in the VOR phase over the training (**Fig. 6h**), nor in the increased OKR gain following the phase reversal training (**Fig. 6f**, compare to **Supplementary Fig. 12a**).

To determine whether the differential activity of TRPC3 ultimately also affects the behavior of the animal, we subjected mice to a task linked to Z– modules, i.e. delay eyeblink conditioning. Mice were trained using a light pulse with 250 ms duration as the conditioned stimulus (CS) and a puff to the cornea as a short unconditioned stimulus (US) at the end of the CS, which over the period of several days evoked conditioned responses (CR, preventative eyelid closure) in the absence of the US (**Fig. 7b**). In contrast to VOR adaptation, the L7-TPRC3^KO^ mice showed significant deficits in eyeblink conditioning during the first week of training (**Fig. 7c**). However, when we subjected them to longer periods they reached similar CR percentages, amplitudes and timing (**Fig. 7d, Supplementary Fig. 12b**).

Thus, although TRPC3 is expressed in both regions underlying the cerebellum-dependent behavioral experiments tested here, TRPC3 activity is selectively required to optimize the cerebellum-dependent learning behavior that is processed in a Z– module^17^. This indicates that the cellular heterogeneity and consequential differentiation in cellular activity also affects the behavior of the animals.

## Discussion

The cerebellum offers a rich repertoire of electrophysiological properties that allows us to coordinate a wide variety of sensorimotor and cognitive behaviors. We recently uncovered that there are probably two main heterogeneous types of cerebellar modules with different intrinsic profiles and plasticity rules^10^. This organization is highly preserved throughout phylogeny and characterized by a series of molecular markers such as zebrin that are distributed in a complementary fashion across the cerebellar cortex^16,41,42^. Here, we demonstrated that zebrin-negative PCs show a relatively high expression of TRPC3, which has a dominant impact on its electrophysiological features. Indeed, gain-of-function and loss-of-function mutations in the gene encoding for TRPC3 selectively affected activity in the zebrin-negative modules and the motor behavior that is controlled by these modules.

TRPC channels, which are calcium-permeable upon activation by phospholipase C or diacylglycerol, are widely expressed in the brain and critically involved in the development and maintenance of synaptic transmission^28,43-45^. TRPC1 and TRPC3 are both prominently expressed in the cerebellum, but in PCs TRPC3 is most abundant^28^. In addition to its contribution to intrinsic activity, TRPC3 currents also mediate the slow excitatory postsynaptic potential following activation of mGluR1b, which is expressed in a pattern complementary to that of zebrin^26,27,45^. The finding that TRPC3 can be detected in all PCs, but that effects of ablation are restricted to zebrin-negative PCs suggests that it is in fact the ‘molecular machinery’ involving mGluR1b activation that drives the differential effects of TRPC3 activation.

In contrast to mGluR1b, mGluR1a is expressed by all PCs (estimated ratio 2:1 to mGluR1b)^27^. The metabotropic receptor mGluR1a is important for IP3-mediated calcium release, climbing fiber elimination as well as PF-PC LTD^26^. Intriguingly, and in line with the concept of modular differentiation, mGluR1-dependent processes are hampered in zebrin-positive PCs by the expression of EAAT4^21^, whereas zebrin-negative PCs selectively express PLCβ4 that works in concert with mGluR1a^26^. The differences in expression patterns may enhance the probability of PF-PC LTD in zebrin-negative PCs over that in zebrin-positive PCs, which is supported by experiments performed in P21 mice^21^. The consequences of EAAT4 or PLCβ4 deletion on PC physiology have been evaluated *in vitro* in several studies^21-24^, but what the consequences *in vivo* on circuit physiology and on the behaviors tested here are, is unclear. Our results here demonstrate that changes that occur at the cell physiological level, i.e. reduced simple spike rate and altered CS-SS interaction, lead to a more complex pattern of changes in the intact system. The additional effects are particularly striking in the L7-TPRC3^KO^ mice, where the reduced simple spike rate in zebrin-negative PCs leads to a lower complex spike rate. In principle, this could have been a direct effect, as lower simple spike rate results in reduced inhibition of the also inhibitory projection from the cerebellar nuclei to the inferior olive^7,46^. However, the unaltered complex spike rate of L7-TRPC3^cKO^ mice suggests that the changes occur during development.

To test the functional consequences of the loss of TRPC3 and the modular specificity of these effects, we tested the impact on behavioral experiments that can be linked to specific modules. Eyeblink conditioning and VOR adaptation are controlled by different modules in the cerebellum and they are distinctly different by nature. Eyeblink conditioning requires a novel, well-timed eyelid movement to a previously unrelated, neutral stimulus, and has been linked to largely or completely zebrin-negative modules in the anterior cerebellum^30,31^. The activity of the putative zebrin-negative PCs in this area is relatively high at rest^47^, in line with their zebrin identity, and a decrease in this high firing rate correlates to the eyeblink response^47-49^. Conversely, VOR adaptation adjusts the amplitude of an existing reflex to optimize sensory processing using visual feedback and is controlled by the vestibulocerebellum, the flocculus in particular, which is classically considered to be zebrin-positive^10,29^ (cf ref.^15,50^). There are more variations in VOR adaptation and the underlying activity patterns are less well-described. In unidirectional VOR gain increase, we recently found that the change correlating with the adapted eye movement consisted of a potentiation, an increase, of the -at rest-lower PC firing rate^51^. Although our current study has its main focus on the differential contribution of TRPC3 at the cell and systems physiological level, it is tempting to speculate how the loss of TRPC3 in PCs results in an eyeblink conditioning phenotype without affecting VOR adaptation. The reduction in firing rate of zebrin-negative PCs may directly contribute to the impaired conditioning. The suppression of simple spike firing that correlates with the conditioned response could be occluded by the lower resting rate in L7-TPRC3^KO^ mice. Alternatively, PF-PC LTD could play a role as it is in line with the simple spike suppression and blocking TRPC3 function completely abolishes this form of LTD^52^. However, genetically ablating PF-PC LTD did not affect the ability to perform eyeblink conditioning successfully^53^, arguing against an exclusive role for this form of plasticity. Schreurs and colleagues demonstrated that intrinsic excitability is increased after eyeblink conditioning^54^. A third option could be that TRPC3 also affects the adaptive increase of excitability, intrinsic plasticity, which is calcium-activated potassium channel function dependent^55^, and thereby delays the expression of a conditioned blink response. All three options would not necessarily affect VOR adaptation and could contribute to the deficits in eyeblink conditioning, but given the relatively mild phenotype, one or two could be sufficient. Future experiments will have to unravel the cellular changes underlying eyeblink conditioning and VOR adaptation and the specific role of TRPC3 in the former.

In this study we aimed to gain insight in the mechanisms that convert molecular heterogeneity into differentiation of cell physiology and function. This mechanistic question goes hand in hand with the more conceptual question: why are there, at least, two different types of PCs? An appealing hypothesis is that zebrin-negative and zebrin-positive bands control two muscles with opposing functions, e.g. a flexor and an extensor. However, trans-synaptic retrograde tracing using rabies virus from antagonist muscles demonstrated that although 3^rd^ order labeling can be found in different parasagittal strips of PCs, there is no apparent division in zebrin-negative and zebrin-positive strips^56^. A second possibility would be that individual muscles are controlled by either only zebrin-negative or zebrin-positive strips, or a combination of both, when needed. In the vestibulocerebellum of the pigeon, each movement direction is controlled by a set of zebrin-negative and zebrin-positive bands^16^. In this configuration each PC within the set, or separately, would then serve a distinct function, for which it is optimized by gene expression patterns. This dissociation of function could entail e.g. timing versus coordination^57^ or moving versus holding still^58^, although none of these distinctions have been linked to specific zebrin-identified modules. Alternatively, it may the net polarity of the connectivity downstream of the cerebellar nuclei up to the motor neurons or the cerebral cortical neurons that determines the demand(s) of the module(s) involved^17^. Module-specific driver lines would greatly aid to answer these questions, but are currently not available.

To summarize, our results support the hypothesis that cerebellar modules control distinct behaviors based on cellular heterogeneity, with differential molecular configurations. We present the first evidence for a non-uniform expression pattern of TRPC3 in PCs, complementary to that of zebrin in the vermis but more homogeneous in the hemispheres. Nonetheless, TRPC3 effects are directly coupled to zebrin, a specificity that putatively requires mGluR1b^26^, the activator of TRPC3 that is expressed in a pattern perfectly complementary to zebrin^27^.

Since the discovery of protein expression patterns in the cerebellar cortex^12^, numerous other proteins with patterned expression have been identified^20^. These patterns have been linked to circuit organizations of modules^41^, to disease and degeneration^20^, and more recently to electrophysiological differences^10,21^. Altogether, this work demonstrates that proper cerebellar function is based on the presence of (at least) two *modi operandi* that have distinct molecular machineries, with a central role for TRPC3, to differentially control sensorimotor integration in downstream circuitries that require control with opposite polarity.

## Materials and Methods

### Animals

For all experiments, we used adult male and female mice with a C57Bl/6 background that were, unless stated otherwise, individually housed, had food ab libitum and were on a 12:12 light/dark cycle. In all experiments the experimenters were blind to mouse genotypes. All experiments were approved by the Dutch Ethical Committee for animal experiments and were in accordance with the Institutional Animal Care and Use Committee.

The generation of TRPC3^Mwk^ mice has been described previously^32^. Briefly, male BALB/cAnN mice carrying the *Mwk* mutation which was generated in a large-scale ENU mutagenesis program were subjected to cross with normal C3H/HeH females, and the first filial generation (F_1_) progeny were screened for a variety of defects. The *Mwk* colony was maintained by repeated backcrossing to C3H/HeH. Experimental mice were generated by crossing C3H/HeH mice heterozygous for the *Mwk* mutation with C57Bl/6 mice. Offspring with the *Mwk* mutation on one allele were classified as gain-of-function TRPC3 Moonwalker mutant (referred to as TRPC3^Mwk^) and littermate mice lacking the *Mwk* mutation were used as controls. Note that, the TRPC3^Mwk^ mutants present evident ataxic phenotype from a very early age, concomitant with progressive degeneration of UBCs and PCs^40^.

Mice in which exon 7 of the TRPC3 gene was flanked by *loxP* sites (TRPC3^fl/fl^ mice)^3^ were bred with mice that express the Cre gene under L7 promoter (PCP2^Cre/-^ mice)^33^. The resulting offspring was genotyped using PCR of genomic DNA extracted from tail or toe by standard procedures. The F_1_ was crossed again with the TRPC3^fl/fl^ mice. Among the second filial generation (F_2_), mice homozygous for the *loxP* sites and one Cre allele were classified as PC-specific TRPC3 knockout (L7^Cre/-^;TRPC3^fl/fl^, here referred to as L7-TRPC3^KO^) mice and as controls when Cre was absent from the genome (L7^−/-^;TRPC3^fl/fl^, here “littermate controls”).

L7-TRPC3^KO^-EAAT4^eGFP^ mice were generated by crossing L7^Cre/-^;TRPC3^fl/fl^ mice with heterozygous EAAT^eGFP/-^ mice which express enhanced green fluorescent protein (eGFP) under control of EAAT4 promotor (**Supplementary Fig. 8**). The F_2_ offspring those who expressed TRPC3^fl/-^, L7^Cre/-^ and EAAT^eGFP/-^ were crossed again with the TRPC3^fl/fl^ mice. Among the F_3_, mice with a homozygous expression of floxed-TRPC3, one Cre allele and one EAAT^eGFP^ allele (L7^Cre/-^;TRPC3^fl/fl^;EAAT4^eGFP/-^), were used and referred to as L7-TRPC3^KO^-EAAT4^eGFP^ mutant mice and as controls when Cre was absent from the genome (L7^−/-^;TRPC3^fl/fl^;EAAT4^eGFP/-^).

Inducible PC-specific TRPC3 knockouts (TRPC3^cKO^) were generated by crossbreeding mice carrying the floxed TRPC3 with mice expressing the tamoxifen-sensitive Cre recombinase Cre-ERT2 under the control of the L7 promoter (obtained from the Institut Clinique de la Souris, www.ics-mci.fr) (experimental mice: L7^Cre-ERT2/-^;TRPC3^fl/fl^). Tamoxifen was dissolved in corn oil to obtain a 20 mg/ml solution, and intraperitoneally injected into all subjects for consecutive 5 days, four weeks prior to electrophysiological recordings. Injections were performed in adults between 12-31 weeks of age. Experimental cohorts were always injected at the same time. Mice without L7^Cre-ERT2^ expression were as control in this study (experimental mice: L7^−/-^;TRPC3^fl/fl^).

### Immunohistochemistry

Anesthetized mice were perfused with 4% paraformaldehyde in 0.12M phosphate buffer (PB). Brains were taken out and post-fixed for 1 h in 4% PFA at room temperature, then transferred in 10% sucrose overnight at 4°C. The next day, the solution was changed for 30% sucrose and left overnight at 4°C. Non-embedded brains were sectioned either sagittally or transversally at 40µm thickness with freezing microtome. Free-floating sections were rinsed with 0.1M PB and incubated 2h in 10mM sodium citrate at 80°C for 2 h, for antigen retrieval. For immuno-fluorescence, sections were rinsed with 0.1M PB, followed by 30 minutes in Phosphate Buffered saline (PBS). Sections were incubated 90 minutes at room temperature in a solution of PBS/0.5%Triton-X100/10% normal horse serum to block nonspecific protein-binding sites, and incubated 48 h at 4°C in a solution of PBS/0.4% Triton-X100/2% normal horse serum, with primary antibodies as follows: Aldolase C (1:500, goat polyclona, SC-12065), Calbindin (1:7000, mouse monoclonal, Sigma, #C9848), and TRPC3 (1:500, rabbit polyclonal, Cell Signaling, #77934). After rinsing in PBS, sections were incubated 2 h at room temperature in PBS/0.4% Triton-X100/2% normal horse serum solution with secondary antibodies coupled with Alexa488, Cy3 or Cy5 (Jackson ImmunoResearch), at a concentration of 1:200. Sections were mounted on coverslip in chrome alum (gelatin/chromate) and covered with Mowiol (Polysciences Inc.). For Light Microscopy section were pre-treated for endogenous peroxidase activity blocking with 3%H_2_O_2_ in PBS, then rinsed for 30 minutes in PBS, incubated 90 minutes in a solution of PBS/0.5%Triton-X100/10% normal horse serum to block nonspecific protein-binding sites, followed by the primary antibody incubation as described before. After 48 h, sections were rinsed in PBS and incubated 2h at room temperature in PBS/0.4% Triton-X100/10% normal horse serum solution with HRP coupled secondary antibodies (Jackson ImmunoResearch), at a concentration of 1:200. Sections were rinsed with 0.1M PB and incubated in diaminobenzidine (DAB, 75 mg/100ml) for 10 minutes. Sections were mounted on glasses in chrome alum (gelatin/chromate), dried with successive Ethanol steps, incubated in Xylene and covered with Permount mounting medium (Fisher Chemical). Images were acquired with an upright LSM 700 confocal microscope (Zeiss) for fluorescent microscopy, and Nanozoomer (Hamamatsu) for light microscopy.

### iDISCO and light sheet imaging

This protocol has been adapted from a previous study^59^. After normal perfusion and post-fixation, brains were washed successively in PBS (1.5 h), 20% methanol/H_2_O (1 h), 50% methanol/H_2_O (1 h), 80% methanol/H_2_O (1 h), and 100% methanol (1 h) twice. To increase clearance, samples were treated with a solution of dichloromethane (DCM) and 100% methanol (2:1) for another hour. Brains were then bleached with 5% H_2_O_2_ in 90% methanol (ice cold) at 4°C overnight. After bleaching, samples successively washed in 80% methanol/H_2_O, 50% methanol/H_2_O, 40% methanol/PBS, and 20% methanol/PBS, for 1 h each, and finally in PBS/0.2% Triton X-100 for 1 h twice. After rehydration, samples were pre-treated in a solution of PBS/0.2% Triton X-100/20% DMSO/0.3 M glycine at 37°C for 36 h, then blocked in a mixture of PBS/0.2% Triton X-100/10% DMSO/6% donkey serum at 37°C for 48 h. Brains were incubated in primary antibody in PTwH solution (PBS/0.2% Tween-20/5% DMSO/3% donkey serum with 10 mg/ml heparin) for 7 days at 37°C with primary antibody: TRPC3 rabbit polyclonal, 1:500 (Cell Signaling, #77934). Amphotericin was added once every two days at 1µg/ml to avoid bacterial growth. Samples were then washed in 24 h in PTwH for six times (1h for each, after the fourth wash, leave it at room temperature overnight), followed by the second round of 7-day incubation with primary antibody. Brains were then washed in PTwH, 6 washes in 24 h, as described before, then incubated in secondary antibody in PTwH/ 3% donkey serum at 37°C for 7 days with secondary anti-Rabbit Cy3 (Jackson ImmunoResearch) at 1:750. Brains were then washed in PTwH, 6 washes in 24 h, again, followed by successive washes in 20% methanol/H_2_O, 40% methanol/H_2_O, 60% methanol/H_2_O, 80% methanol/H_2_O, and 100% methanol twice, for 1 h each, and finally incubation overnight in a solution of DCM and100% methanol. For tissue clearing, brains were incubated 20 mins in DCM, twice, and conserved in Benzyl ether at room temperature.

Ready samples were imaged in horizontal orientation with an UltraMicroscope II (LaVision BioTec) light sheet microscope equipped with Imspector (version 5.0285.0) software (LaVision BioTec). Images were taken with a Neo sCMOS camera (Andor) (2560×2160 pixels. Pixel size: 6.5 x 6.5 μm2). Samples were scanned with double-sided illumination, a sheet NA of 0.148348 (resuls in a 5 μm thick sheet) and a step-size of 2.5 μm using the horizontal focusing light sheet scanning method with the optimal amount of steps and using the contrast blending algorithm. The effective magnification for all images was 1.36x (zoombody*objective + dipping lens = 0.63x*2.152x). Following laser filter combinations were used: Coherent OBIS 488-50 LX Laser with 525/50nm filter, Coherent OBIS 561-100 LS Laser with 615/40 filter, Coherent OBIS 647-120 LX with 676/29 filter.

### Western blot and fractionation

Cerebellar tissue from L7-TRPC3^KO^ and control mice was dissected and immediately frozen in liquid nitrogen. Samples were homogenized with a Dounce homogenizer in lysis buffer containing 50 mM Tris-HCl pH 8, 150 mM NaCl, 1% Triton X-100, 0.5% sodium deoxycholate, 0.1% SDS and protease inhibitor cocktail. Protein concentrations were measured using Pierce BCA protein assay kit (Thermo Fisher). Samples were denatured and proteins were separated by SDS-PAGE in Criterion™ TGX Stain-Free™ Gels (Bio-Rad), and transferred onto nitrocellulose membranes with the Trans-Blot^®^ Turbo™ Blotting System (Bio-Rad). Membranes were blocked with 5% BSA (Sigma-Aldrich) in TBST (20mM Tris-HCl pH7.5, 150mM NaCl and 0.1%, Tween20) for 1 h and probed with the following primary antibodies: rabbit anti-TRPC3 (Cell Signaling Technology, #77934; 1:1000) and mouse anti-actin (Millipore, MAB1501; 1:1000). Secondary antibodies used were IRDye 800CW Donkey anti-Rabbit IgG (LI-COR Biosciences, Cat # 925-32213; 1:20000) and IRDye 680RD Donkey anti-Mouse IgG (LI-COR Biosciences, Cat # 925-68072; 1:20000). Membranes were scanned by Odyssey Imager (LI-COR Biosciences) and quantified using Image Studio Lite (LI-COR Biosciences). For quantification, densitometry of protein bands of interest was normalized to that of actin.

For fractionation experiments, cerebellar tissues from C57/BL6 were collected and the synaptosomes were isolated using Syn-PER^TM^ Synaptic Protein Extraction Reagent (ThermoScientific, #87793) according to the manufacturer’s instructions.

### *In vivo* extracellular recordings and analysis

We performed in vivo extracellular recordings in adult TRPC3^Mwk^ (aged 15-47 weeks), L7-TRPC3^KO^ (aged 22-43 weeks), L7-TRPC3^cKO^ (aged 17-28 weeks) mice, respectively, as previously described^5^. Briefly, an immobilizing pedestal consisting of a brass holder with a neodymium magnet (4×4×2 mm) was fixed on the skull, overlying the frontal and parietal bones, and then a craniotomy (Ø 3 mm) was made in the interparietal or occipital bone under general anesthesia with isoflurane/O_2_ (4% induction, 1.5-2% maintenance). After over 24 h of recovery, mice were head-fixed and body restrained for recordings. PCs were recorded from vermal lobules I-III and X, using a glass pipette (OD 1.5 mm, ID 0.86 mm, borosilicate, Sutter Instruments, USA; 1-2 µm tips, 4-8 MΩ) with a downward pitch angle of 40° and 65° respectively. The pipettes were filled with 2 M NaCl-solution and mounted on a digital 3-axis drive (SM-5, Luigs Neumann, Germany). After recording, biotinylated dextran amines (BDA) was iontophoretically injected to confirm that the recordings were from Lobules I-III or X. PCs were identified by the presence of simple and complex spikes, and determined to be from a single unit by confirming that each complex spike was followed by a climbing fiber pause. All in vivo recordings were analyzed offline using Spiketrain (Neurasmus BV, Rotterdam, The Netherlands), running under MatLab (Mathworks, MA, USA). For each cell, the firing rate, CV and mean CV2 were determined for simple and complex spikes, as well as the climbing fiber pause. The CV is calculated by dividing the standard deviation, SD, by the mean of ISIs, whereas CV2 is calculated as 2×|ISI_n+1_-ISI_n_| / (ISI_n+1_+ISI_n_). Both are measures for the regularity of the firing, with CV reflecting that of the entire recording and mean CV2 that of adjacent intervals, making the latter a measure of regularity on small timescales. The climbing fiber pause is determined as the duration between a complex spike and the fist following simple spike. To extend this analysis, we also plotted histograms of simple spike activity time locked on the complex spike, and labelled the shape of this time histogram as normal, facilitation, suppression, or oscillation^5^.

### *In vivo* two-photon targeted electrophysiology

Details on targeted electrophysiological recordings in vivo in the mouse cerebellum were described previously^36^. PCs in lobules IV-VI were recorded in adult L7-TRPC3^KO^-EAAT4^eGFP^ mice (aged 14-49 weeks) under two-photon microscope guidance. A custom-made head plate was fixed to the cleaned skull of each animal, under isoflurane anesthesia, with dental adhesive (Optibond; Kerr corporation, West collins, USA) and secured with dental acrylic. A craniotomy was made above the cerebellum, exposing lobules IV-VI. The craniotomy was sealed with biocompatible silicone (Kwik-Cast; World Precision Instruments) and the animal was allowed to recover from surgery before recording. The silicone seal was removed prior to recording. To keep the brain surface moist, Ringer solution containing (in mM): NaCl 135, KCl 5.4, MgCl_2_ 1, CaCl_2_ 1.8, HEPES 5 (pH 7.2 with NaOH; Merck, Darmstadt, Germany) was applied. Glass micropipettes with tip size of ~1 μm (resistance: 6-9 MΩ) were advanced from the dorsal surface under a 25° angle into the cerebellum, allowing concurrent two-photon imaging with a long working distance objective (LUMPlanFI/IR 40×/0.8; Olympus) on a custom-built two-photon microscope. Pipettes were filled with the same Ringer solution with an additional 40 μM AlexaFluor 594 hydrazide (Sigma-Aldrich, Steinheim, Germany) for visualization. GFP and AlexaFluor 594 were simultaneously excited by a MaiTai laser (Spectra Physics Lasers, Mountain View, CA, USA) operated at 860 nm. Green (GFP) and red (AlexaFluor 594) fluorescence were separated by a dichroic mirror at 560nm and emission filters centered at 510nm (Brightline Fluorescence Filter 510/84; Semrock) and 630nm (D630/60; Chroma), respectively. The brain surface was stabilized with agarose (2% in Ringer; Sigma–Aldrich) and pipette pressure was initially kept at 3 kPa while entering the brain tissue. It was then removed for cell approach and the actual recording. Extracellular potentials were acquired with a MultiClamp 700A amplifier (Molecular Devices, Sunnyvale, CA, USA) in current-clamp mode. Signals were low-pass filtered at 10 kHz (four-pole Bessel filter) and digitized at 25 kHz (Digidata 1322A). Data were recorded with pCLAMP 9.2 (Molecular Devices). Z+ and Z– cells were identified by comparing the relative intensity of GFP fluorescence. Whenever possible, cells of both types were recording alternatingly between adjacent bands Purkinje neurons with high and low GFP fluorescence.

### *In vitro* electrophysiology and analysis

We performed *in vitro* electrophysiological recordings on TRPC3^Mwk^ (aged 9-21 weeks) and L7-TRPC3^KO^ (aged 20-60 weeks). As described previously^60^, acute sagittal slices (250 μm thick) were prepared from the cerebellar vermis and put into ice-cold slicing medium which contained (in mM) 240 sucrose, 2.5 KCl, 1.25 Na_2_HPO_4_, 2 MgSO_4_, 1 CaCl_2_, 26 NaHCO_3_ and 10 D-Glucose, carbogenated continuously with 95% O_2_ and 5% CO_2_. After cutting using a vibrotome (VT1200S, Leica), slices were incubated in artificial cerebrospinal fluid (ACSF) containing (in mM): 124 NaCl, 5 KCl, 1.25 Na_2_HPO_4_, 2 MgSO_4_, 2 CaCl_2_, 26 NaHCO_3_ and 15 D-Glucose, equilibrated with 95% O_2_ and 5% CO_2_ at 33.0±1.0 °C for 30 min, and then at room temperature. NBQX (10 μM), DL-AP5 (50 μM), and picrotoxin (100 μM) were bath-applied to block AMPA-, NMDA-, and GABA subtype A (GABA_A_)-receptors, respectively. PCs were identified using visual guidance by DIC video microscopy and water-immersion 40X objective (Axioskop 2 FS plus; Carl Zeiss, Jena, Germany). Recording electrodes (3-5 MΩ, 1.65 mm outside diameter and 1.11 mm interior diameter (World Precision Instruments, Sarasota, FL, USA) were prepared using a P-97 micropipette puller (Sutter Instruments, Novato, CA, USA), and filled with ACSF for cell-attached recordings, or with an intracellular solution containing (in mM): 120 K-Gluconate, 9 KCl, 10 KOH, 4 NaCl, 10 HEPES, 28.5 Sucrose, 4 Na_2_ATP, 0.4 Na_3_GTP (pH 7.25-7.35 with an osmolality of 295) for whole-cell recordings. We measured spontaneous firing activity of PCs in cell-attached mode (0 pA injection) and intrinsic excitability in whole-cell current-clamp mode by injection of brief (1s) depolarizing current pulses (ranging from −100 to 1100pA with 100pA increments) from a membrane holding potential of –65 mV at 33.0±1.0°C. The spike count of evoked action potential was taken as a measure of excitability. AP properties including peak amplitude, after-hyperpolarization amplitude (AHP) and half-width were evaluated using the first action potential generated by each PC. AHP indicates the amplitude of undershoot relative to the resting membrane potential. Half-width indicates the width of the signal at 50% of the maximum amplitude. PCs that required > −800 pA to maintain the holding potential at −65 mV or fired action potentials at this holding potential were discarded. The average spiking rate measured over the entire current pulse was used to construct current-frequency plots. For whole-cell Recordings, cells were excluded if the series (Rs) or input resistances (Ri) changed by >15% during the experiment, which was determined using a hyperpolarizing voltage step relative to the −65 mV holding potential. All electrophysiological recordings were acquired in lobules I-III and lobule X of the vermal cerebellum using EPC9 and EPC10-USB amplifiers (HEKA Electronics, Lambrecht, Germany) and Patchmaster software (HEKA Electronics). Data were analyzed afterwards using Clampfit (Molecular Devices).

### Compensatory eye movement recordings

We subjected alert L7-TRPC3^KO^ mice (aged 12-39 weeks) to compensatory eye movement recordings which were described in detail previously^61^. In short, mice were equipped with a pedestal under general anesthesia with isoflurane/O_2_. After a 2-3 days of recovery, mice were head-fixed with the body loosely restrained in a custom-made restrainer and placed in the center of a turntable (diameter: 63 cm) in the experimental set-up. A round screen (diameter 60 cm) with a random dotted pattern (‘drum’) surrounded the mouse during the experiment. Compensatory eye movements were induced by sinusoidal rotation of the drum in light (OKR), rotation of the table in the dark (VOR) or the rotation of the table in the light (visually enhanced VOR, VVOR) with an amplitude of 5° at 0.1-1 Hz. Motor performance in response to these stimulations was evaluated by calculating the gain (eye velocity/stimulus velocity) and phase (eye to stimulus in degrees) of the response. Motor learning was studied by subjecting mice to mismatched visual and vestibular input. Rotating the drum (visual) and table (vestibular) simultaneously, in phase at 0.6 Hz (both with an amplitude of 5°, 5 x 10 min) in the light will induce an increase of the gain of the VOR (in the dark). Subsequently, VOR Phase reversal was tested by continuing the next days (day 2-5, keeping mice in the dark in between experiments) with in phase stimulation, but now with drum amplitudes of 7.5° (days 2) and 10° (days 3, 4, and 5), while the amplitude of the turntable remained 5°. This resulted, over days of training, in the reversal of the VOR direction, from a normal compensatory rightward eye movement (in the dark), when the head turns left, to a reversed response with a leftward eye movement, when the head moves left. At the end of the VOR phase reversal training the OKR was probed again and compared to the OKR before training, to examine OKR gain increase. VOR gain increase was evoked by subjecting mice to out of phase drum and table stimulation at 1.0 Hz (both with an amplitude of 1.6°). A CCD camera was fixed to the turntable in order to monitor the eyes of the mice. Eye movements were recorded with eye-tracking software (ETL-200, ISCAN systems, Burlington, NA, USA). For eye illumination during the experiments, two infrared emitters (output 600 mW, dispersion angle 7°, peak wavelength 880 nm) were fixed to the table and a third emitter, which produced the tracked corneal reflection, was mounted to the camera and aligned horizontally with the optical axis of the camera. Eye movements were calibrated by moving the camera left-right (peak-to-peak 20°) during periods that the eye did not move^62^. Gain and phase values of eye movements were calculated using custom-made Matlab routines (MathWorks).

### Eyeblink conditioning

For all procedures on eyeblink conditioning we refer to the study done previously^63^. L7-TRPC3^KO^ mice, aged 16-25 weeks, were anesthetized with an isoflurane/oxygen mixture and surgically placed a so-called pedestal on the skull. After a 2-3 days’ recovery, mice were head-fixed and suspended over a foam cylindrical treadmill on which they were allowed to walk freely (**Fig. 7b**). Before each session starting, a minuscule magnet (1.5×0.7×0.5mm) was placed on the left lower eyelid with superglue (cyanoacrylate) and an NVE GMR magnetometer was positioned above the left upper eyelid. With this magnetic distance measurement technique (MDMT), we measured the exact positions of each individual mouse eyelid by analyzing the range from optimal closure to complete aperture. The CS was a green LED light (CS duration 280 ms, LED diameter 5 mm) placed 10 cm in front of the mouse’s head. The US consisted of a weak air-puff applied to the eye (30 psi, 30 ms duration), which was controlled by an API MPPI-3 pressure injector, and delivered via a 27.5-gauge needle that was perpendicularly positioned at 0.5-1 cm from the center of the left cornea. The training consisted of 3 daily habituation sessions, 1 baseline measurement, 3 blocks of 5 daily acquisition sessions (each block was separated by 2 days of rest). During the habituation sessions, mice were placed in the setup for 30-45 minutes, during which the air puff needle (for US delivery) and green LED (for CS delivery) were positioned properly but no stimuli were presented. On the day of acquisition session 1, each animal first received 20 CS-only trials as a baseline measure, to establish that the CS did not elicit any reflexive eyelid closure. During each daily acquisition session, every animal received in total 200 paired CS-US trials, 20 US only trials, and 20 CS only trials. These trials were presented in 20 blocks, each block consisted of 1 US only trial, 10 paired CS-US trials, and 1 CS only trial. Trials within the block were randomly distributed, but the CS only trial was always preceded by at least 2 paired CS-US trials. The interval between the onset of CS and that of US was set at 250 ms. All experiments were performed at approximately the same time of day by the same experimenter. Individual eyeblink traces were analyzed automatically with custom computer software (LabVIEW or MATLAB). Trials with significant activity in the 500 ms pre-CS period (>7*IQR) were regarded as invalid for further analysis. Valid trials were further normalized by aligning the 500 ms pre-CS baselines and calibrating the signal so that the size of a full blink was 1. In valid normalized trials, all eyelid movements larger than 0.1 and with a latency to CR onset between 50-250 ms and a latency to CR peak of 100-250 ms (both relative to CS onset) were considered as conditioned responses (CRs). For CS only trials in the probe session we used the exact same criteria except that the latency to CR peak time was set at 100-500 ms after CS onset.

### Erasmus Ladder

Mice aged 11-16 weeks were subjected to the Erasmus Ladder (Noldus, Wageningen, Netherlands). As described previously^9^, the Erasmus Ladder is a fully automated system consisting of a horizontal ladder between two shelter boxes. The ladder has 2 x 37 rungs for the left and right side. Rungs are placed 15 mm apart, with alternate rungs in a descended position, so as to create an alternating stepping pattern with 30 mm gaps. All rungs are equipped with touch sensors, which are activated when subject to a pressure corresponding to more than 4 grams. The sensors are continuously monitored to record the position and the walking pattern of the mouse. A single crossing of the Erasmus Ladder is recorded as a trial. In this study, each mouse underwent a daily session consisting of 42 trials, for five consecutive days. Motor performance was measured by counting step durations and percentages during a trial, including short steps (steps from one high rung to the next high rung), long steps (skipping one high rung), jumps (skipping two high rungs), lower steps (a step forward steps, but the paw is placed on a low rung), back steps (a step backward steps from one high rung to the previous high rung). All data were collected and processed by ErasmusLadder 2.0 software (Noldus, Wageningen, Netherlands).

### Statistics

All values are shown as mean ± s.d., unless stated otherwise. Inter-group comparisons were done by two-tailed Student's t test. For combined analysis of multiple sections, ANOVA for repeated measures was used to analyze eye-movement recording data; linear mixed-effect model analysis^63^ (established in R version 1.1.442) was used to analyze eyeblink conditioning data. For a complete dataset, see **Supplementary Table 2-6**. All statistical analyses were performed using SPSS 20.0 software. Data was considered statistically significant if *P* < 0.05. * *P* < 0.05, ** *P* < 0.01, *** *P* < 0.001.

## Supporting information

## Data availability

Data supporting the findings of this study are available from the corresponding author upon reasonable request.

## Acknowledgements

We kindly thank Laura Post for mouse breeding; Nadia Khosravinia for help with behavior experiment; Joshua J. White and Dick Jaarsma for discussions and comments on the manuscript. This work was supported by an ERC starter grant (ERC-Stg #680235; MS), China Scholarship Council (#201306230130; BW), the Netherlands Organization for Scientific Research (NWO-ALW; CIDZ), the Dutch Organization for Medical Sciences (ZonMW; CIDZ), ERC-adv and ERC-POC (CIDZ), and the Center for Integrated Protein Science Munich (CIPSM; JH).

## Author contributions

B.W. and M.S. designed all the experiments and wrote the manuscript; B.W. performed and analyzed the *in vivo* and *in vitro* electrophysiology experiments, analyzed the *in vivo* two-photon experiments and the eye-movement behavior experiments; F.B. performed and analyzed the immunohistochemistry and iDISCO experiments; A.B.W performed the *in vivo* two-photon experiments; C.O. conducted and analyzed the Western Blot; Y.A. and RJ.P supported for light sheet imaging; J.H. supplied the TRPC3^fl/fl^ mice; E.B. supplied the TRPC3^Mwk^ mouse; HJ.B analyzed the eyeblink conditioning data; C.I.D.Z. provided major revisions to the manuscript and guided the project. M.S. initiated the project and coordinated collaborations between groups. All authors discussed the results and implications and commented on the manuscript.

## Additional information

**Supplementary information** linked to this paper is available on *Nature Communications’* website.

