## Supplementary material for "TRPC3 is essential for functional heterogeneity of cerebellar Purkinje cells"

### Supplementary information

Supplementary information for this article includes 12 figures, 6 tables and 1 movie.

### Supplementary Figure Legends

**Supplementary Fig.1 | Overview of TRPC3 expression pattern, related to Fig.1**

Immunofluorescent images of coronal sections of wild-type mouse cerebellar cortex, from rostral to caudal, stained with anti-TRPC3 (red) and anti-aldolase C (green). TRPC3 is expressed in evident parasagittal bands which are complementary to Aldolase C (left) in the vermis, but more uniform in the hemispheres. For more detailed pictures, see **Supplementary Fig. 2**. Cr II, Crus II; PM, paramedian lobule; Cop, copula of the pyramis; Sim, simple lobule; PFL, paraflocculus; FL, flocculus. Scale bar: 500 µm.

**Supplementary Fig.2 | Local patterns of TRPC3 expression, related to Fig.1**

Immunofluorescent images of coronal sections of wild-type cerebellum, illustrating the distribution of TRPC3 in the anterior vermis (**a**), posterior vermis (**b**) and hemispheres (**c**). Demarcated areas in top row of **a** and **b** are magnified underneath. Note that TRPC3 immunoreactivity is moderately higher in the Z– PCs than that in the Z+ PCs in the vermis. In the hemispheres the TRPC3-labeled bands are less well defined and can either be complementary to (**c**, top) or indistinguishable from (**c**, bottom) zebrin bands. ml, molecular layer; gcl, granule cell layer; pcl, purkinje cell layer; Scale bar: 50 µm in (**a, b**); 100 µm in (**c**).

**Supplementary Fig.3 | Western blot analysis of L7-TRPC3$^{KO}$ mice, related to Fig.2**

**a**, Representative western blots (top) and quantification (bottom) show significantly reduced TRPC3 levels in both anterior (left) and posterior (right) cerebellum in L7-TRPC3$^{KO}$ mice. Note the residual TRPC3 present in the posterior cerebellum, presumably due to the presence of unaffected TRPC3-expressing UBCs, which are virtually absent in the anterior cerebellum. **b**, Images of full-length western blots presented in (**a**). **c**, Schematic for synaptic protein extraction protocol. **d**, Subcellular localizations by western blots in the anterior (top) and posterior (bottom) cerebellum. TRPC3 is abundantly present in the membrane (P1) and synaptosomes (P2), but less so in the cytosol (S2). L7-TRPC3$^{KO}$ mice were devoid of TRPC3 completely in both anterior and posterior cerebellar fractionations. **e**, Images of full-length western blots presented in (**d**).

**Supplementary Fig.4 | Immunohistochemical analysis of L7-TRPC3$^{KO}$ mice, related to Fig.2**

**a-b**, Coronal immunofluorescence images of stainings for TRPC3 (red) and Aldolase C (green) in the posterior cerebellar cortex of L7-TRPC3$^{KO}$ mutants (right) and normal mice (left) with a higher magnifications of the squared areas (right). In contrast to the TPRC3 staining in control mice there is no longer a banding pattern visible for TRPC3 in mutant mice, while aldolase C is still clearly present in bands. The presence of TRPC3 staining in the UBCs, see e.g. the example indicated by the arrowhead, confirms that the antibody worked and that the loss of TRPC3 is specific for PCs (marked by asterisks).

**Supplementary Fig.5 | Cell-attached recordings of PC activity in TRPC3 mutants *in vitro*, related to Fig.2**

**a,** Representative traces of PCs (top) and the corresponding inter spike interval (ISI) distributions of Z– (left) and Z+ (right) PCs of L7-TRPC3$^{KO}$ mice. **b-c**, The coefficient of variation (CV), measure for regularity of the entire trace, and CV2, measure of regularity on short time-scales, of ISIs were both significantly reduced in Z– PCs of L7-TRPC3$^{KO}$ mice (light-green, n=40 cells/N=6 mutants vs. n=43/N=5 controls; CV: $t_{79}$=2.14, $P$=0.036; CV2: $t_{79}$=2.54, $P$=0.013), but unaltered in Z+ PCs (dark-green, n=36/N=10 vs. n=35/N=4, CV: $t_{71}$=-1.13, $P$=0.263; CV2: $t_{67}$=-0.977, $P$=0.332), compared with littermate controls. (**d**) Similar to (**a**) but for TRPC3$^{Mwk}$ mice. **e-f**, PCs of TRPC3$^{Mwk}$ mice showed no significant differences *in vitro* in CV and CV2, either in Z– PCs (light-red, n=15/N=4 mutants vs. n=11/N=2 controls; CV: $t_{24}$=0.34, $P$=0.735; CV2: $t_{24}$=1.32, $P$=0.199), or in Z+ PCs (dark-red, n=13/N=4 vs. n=10/N=2 controls; CV: $t_{21}$=0.985, $P$=0.336; CV2: $t_{21}$=0.960, $P$=0.348), compared with littermate controls. Error bars denote s.d.. Lighter colors represent Z– and darker colors represent Z+ PCs, respectively. See **Supplementary Table 2** for values.

**Supplementary Fig.6 | Quantification of *in vitro* patch-clamp recordings of PC activity in L7-TRPC3$^{KO}$ mice, related to Fig.2**

Whole-cell patch clamp recordings of PCs of L7-TRPC3$^{KO}$ mice, revealed no significant differences in holding current or parameters of first action potential evoked by current injection, including peak-amplitude, half-width and AHP, between mutants and controls. Error bars denote s.e.m. Lighter colors represent Z– and darker colors represent Z+ PCs, respectively. See **Supplementary Table 2** for values and statistics.

**Supplementary Fig.7 | *In vivo* extracellular recordings of PC activity in L7-TRPC3$^{KO}$ and TRPC3$^{Mwk}$ mice, related to Figure 3 and 5**

**a,g**, Representative Purkinje cell recording traces of TRPC3$^{Mwk}$ and L7-TRPC3$^{KO}$ mice, respectively. Asterisks indicate complex spikes, lighter colors represent Z– and darker colors represent Z+ PCs, respectively. **b-f**, CV, a measure for regularity of the entire trace, for simple spikes (SS-CV) as well as for complex spikes (CS-CV) of PCs recorded in TRPC3$^{Mwk}$ mice, were significantly increased in both Z– and Z+ PCs. In Z– PCs this can exclusively be attributed to TRPC3 gain-of-function, while in Z+ PCs the loss of the regular input from UBCs (predominantly present in Z+ areas) potentially contributes to the phenotype. CV2, a measure for regularity on short time-scales, for simple spikes (SS-CV2), as well as for complex spikes (CS-CV2) of PCs recorded, and CF-pause in both Z– and Z+ PCs of TRPC3$^{Mwk}$ mice were unaffected, compared with those of littermate controls. **h-l**, In L7-TRPC3$^{KO}$ mice, SS-CV, SS-CV2, CS-CV, CS-CV2 in both Z– and Z+ PCs do not differ from littermate controls. However, CF-pause was significant longer Z– Purkinje cell (**l**, left), but unaffected in Z+ PCs. Error bars denote s.d.. See **Supplementary Table 3** for values and statistics.

**Supplementary Fig.8 | Breeding strategy to obtain L7-TRPC3$^{KO}$-EAAT4$^{eGFP}$ mice, related to Fig.4**

L7-TRPC3$^{KO}$-EAAT4$^{eGFP}$ mice were generated by first crossing female L7$^{Cre/-}$;TRPC3$^{fl/fl}$ mice with male EAAT4 $^{eGFP/-}$ mice. Heterozygous female knockouts (Het KO, only one allele of TRPC3 gene floxed by *loxp*) were then crossed with homozygous male TRPC3$^{fl/fl}$ mice. The final F3 cross generated littermates that TRPC3$^{fl/fl}$; EAAT4 $^{eGFP/-}$ with no Cre present (controls) and with heterozygous Cre specific in PC neurons (mutants).

**Supplementary Fig.9 | *In vivo* extracellular recordings of PC activity in L7-TRPC3$^{KO}$-EAAT4$^{eGFP}$ and L7-TRPC3$^{cKO}$ mice, related to Fig. 4 and 5**

**a,g**, Representative PC recording traces of L7-TRPC3$^{KO}$-EAAT4$^{eGFP}$ and L7-TRPC3$^{cKO}$ respectively. Asterisks indicate complex spikes, lighter colors represent Z– and darker colors represent Z+ PCs, respectively. **b-f**, In L7-TRPC3$^{KO}$-EAAT4$^{eGFP}$ mice, all parameters for Z– PCs, including SS-CV, SS-CV2, CS-CV, CS-CV2 and CF-pause, were significantly increased compared to those of littermate controls. However, there was no change in those features of the Z+ PCs. **h-l**, PCs in L7-TRPC3$^{cKO}$ mice showed, after tamoxifen-induced TRPC3 ablation, no significant differences in SS-CV, SS-CV2, CS-CV, CS-CV2 and CF-pause in Z– or Z+ PCs, as compared with those of littermate controls that were also injected with tamoxifen. Error bars denote s.d.. See **Supplementary Table 3** for values and statistics.

**Supplementary Fig.10 | L7-TRPC3$^{KO}$ mice show normal Erasmus ladder performance**

Motor coordination measured by Erasmus Ladder did not differ between L7-TRPC3$^{KO}$ mice (N=16) and their wildtype littermates (N=16). **a**, The setup of Erasmus Ladder which consists of a horizontal ladder (magnifications on the right) connecting two shelter boxes. **b**, Schematic of high rungs (green) and low rungs (red) with purple arrows illustrating the five different step types: ① Back steps, ② Short steps, ③ Long steps, ④Jumps, ⑤ Lower steps (see methods). **c**, The distribution of step types in L7-TRPC3$^{KO}$ mice did not differ from their littermate controls over the five days tested. Values are shown as mean±s.e.m., see **Supplementary Table 6** for values and statistics.

**Supplementary Fig.11 | TRPC3 immunoreactivity in areas related to eyeblink conditioning and compensatory eye movements, related to Fig. 6 and 7**

**a**, The region of eyeblink conditioning, locating at the sulcus between lobule IV-V and VI, is largely Z– (middle), and stained positive for TRPC3 (top). **b**, Compensatory eye movements and their adaptation are under control of the flocculus of the cerebellum. Zebrin-staining is more positive in the flocculus, while being less intense for TRPC3. Note that TRPC3 staining is not absent in the flocculus and that UBCs in the granule cell layer of the flocculus also stain positive for TRPC3. ml, molecular layer; gcl, granule cell layer; pcl, purkinje cell layer; prf, primary fissure; Sim, simple lobule; PFL, paraflocculus; FL, flocculus; CO, cochlear nucleus. Scale bars: 200 μm.

**Supplementary Fig.12 | L7-TRPC3$^{KO}$ mice show normal motor performance and timing in flocculus-dependent compensatory eye movements and eyeblink conditioning, related to Fig. 6 and 7**

**a**, Gain (top) and phase (bottom) of baseline performance of compensatory eye movements: the optokinetic reflex (OKR), the vestibulo-ocular reflex (VOR) and the visually-enhanced VOR (VVOR) were not affected in L7-TRPC3$^{KO}$ mice, compared to littermate controls

(N=13 versus N=11, all $P > 0.05$). **b**, Peri-stimulus histogram plots with a Gaussian kernel density estimate (black dashed line) showing the distribution of CR onset (dark filled bars) and CR peak time (light filled bars) relative to CS and US onset in CS only trials for session 1, 5, 10, and 15. In both groups, there is a clear development in CR onset and peak time: there are no clearly preferred times in the CS-US interval at the start of training (session 1), but during training CR onset values are centered around 100-125 ms after CS onset, and CR peak times are located around the onset of the expected US. Green dashed line is CS onset, red dashed line is US onset; light green and light red fill indicate CS and US duration, respectively. **c**, Two-dimensional density plot showing latency to CR peak relative to the fraction eyelid closure over all sessions. Both groups clearly show CRs that are timed around the onset of US. Values are shown as mean±s.e.m., see **Supplementary Table 4-5** for values and statistics.

**Supplementary Movie, related to Fig.1**

Light sheet imaging reconstruction of whole-mount immunolabeling for TRPC3 (white signal), scanned in the horizontal plane of an adult mouse brain, cleared with iDISCO protocol (see Methods).

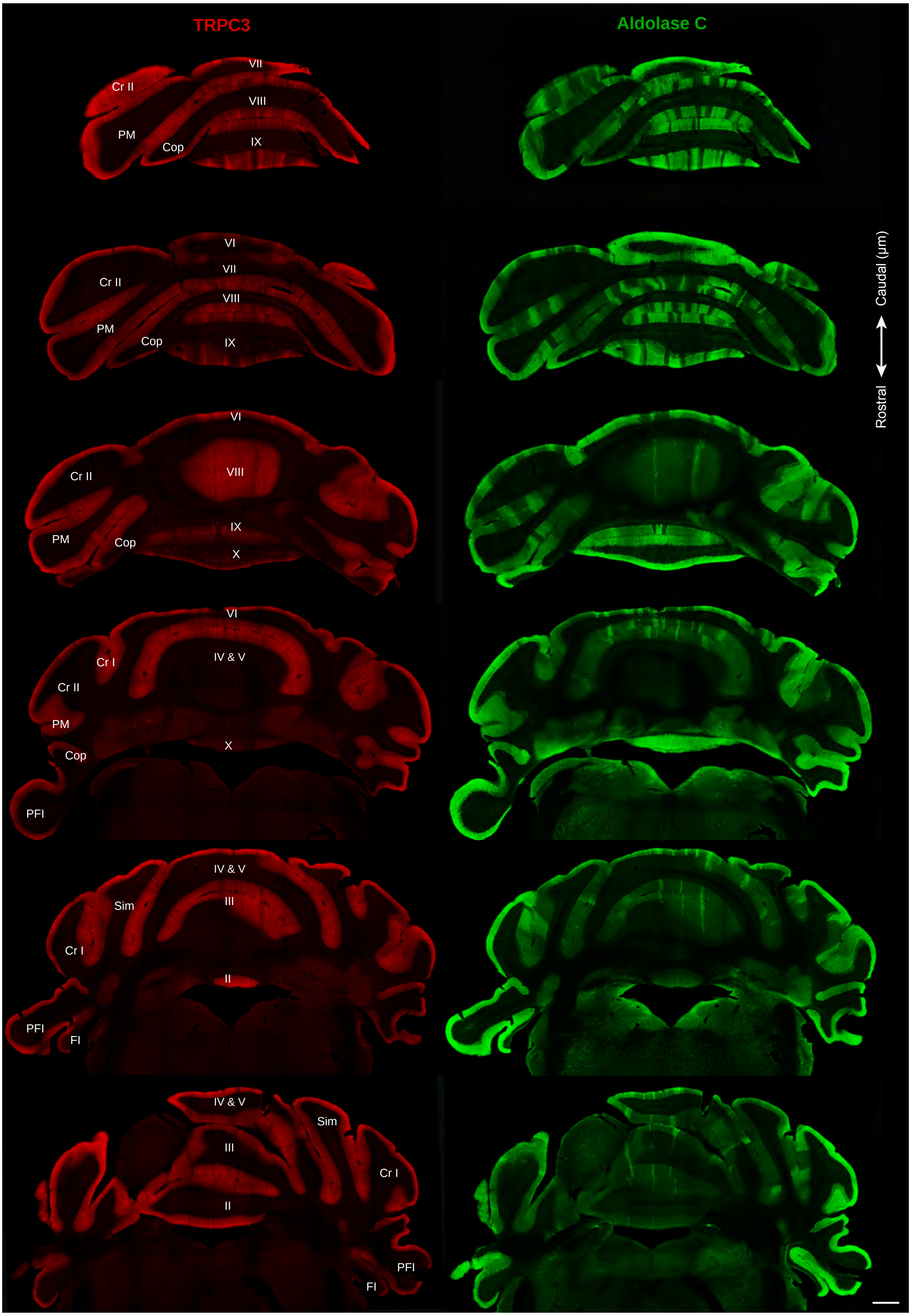


**Supplementary fig.1**

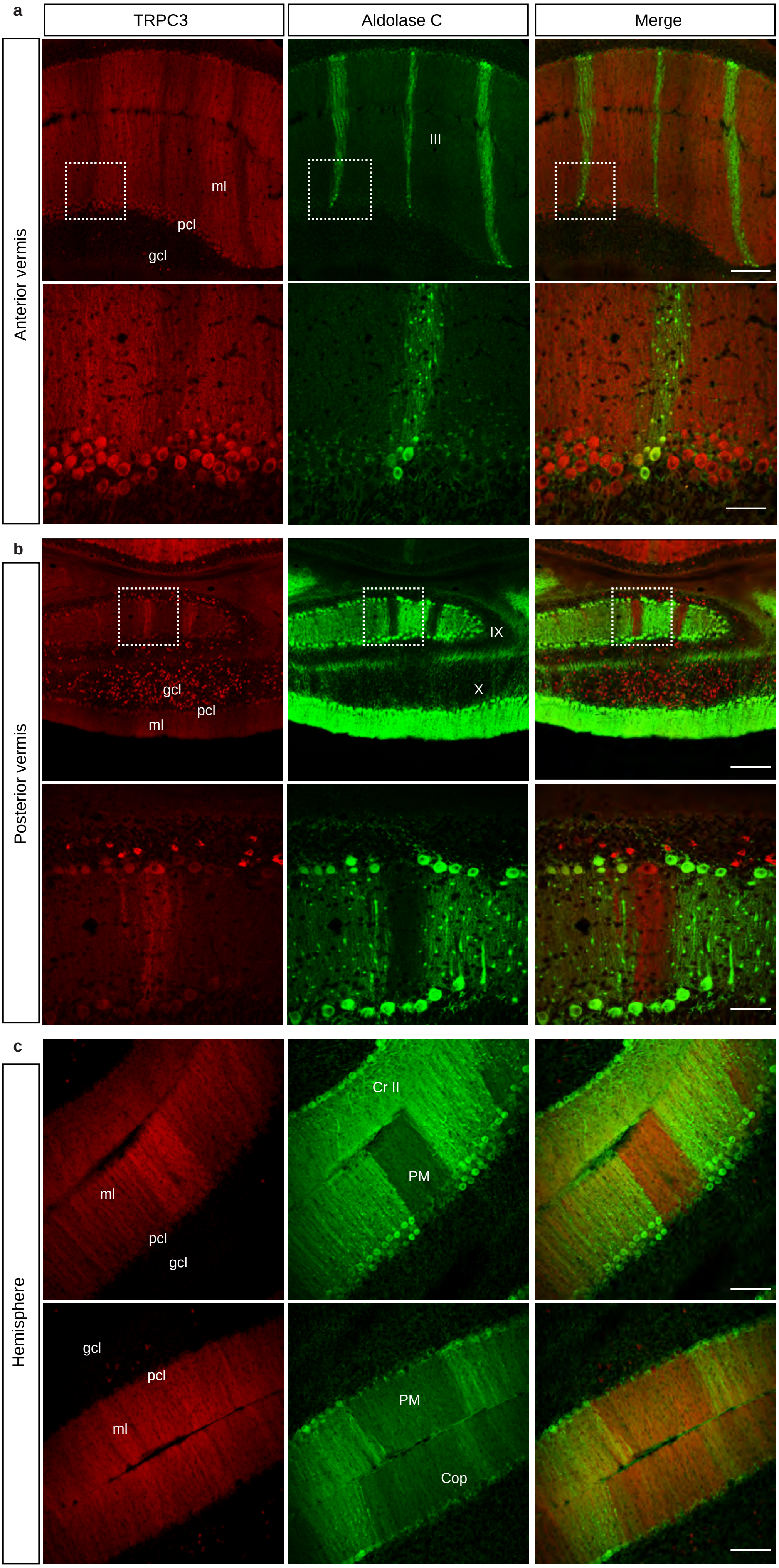


**Supplementary fig.2**

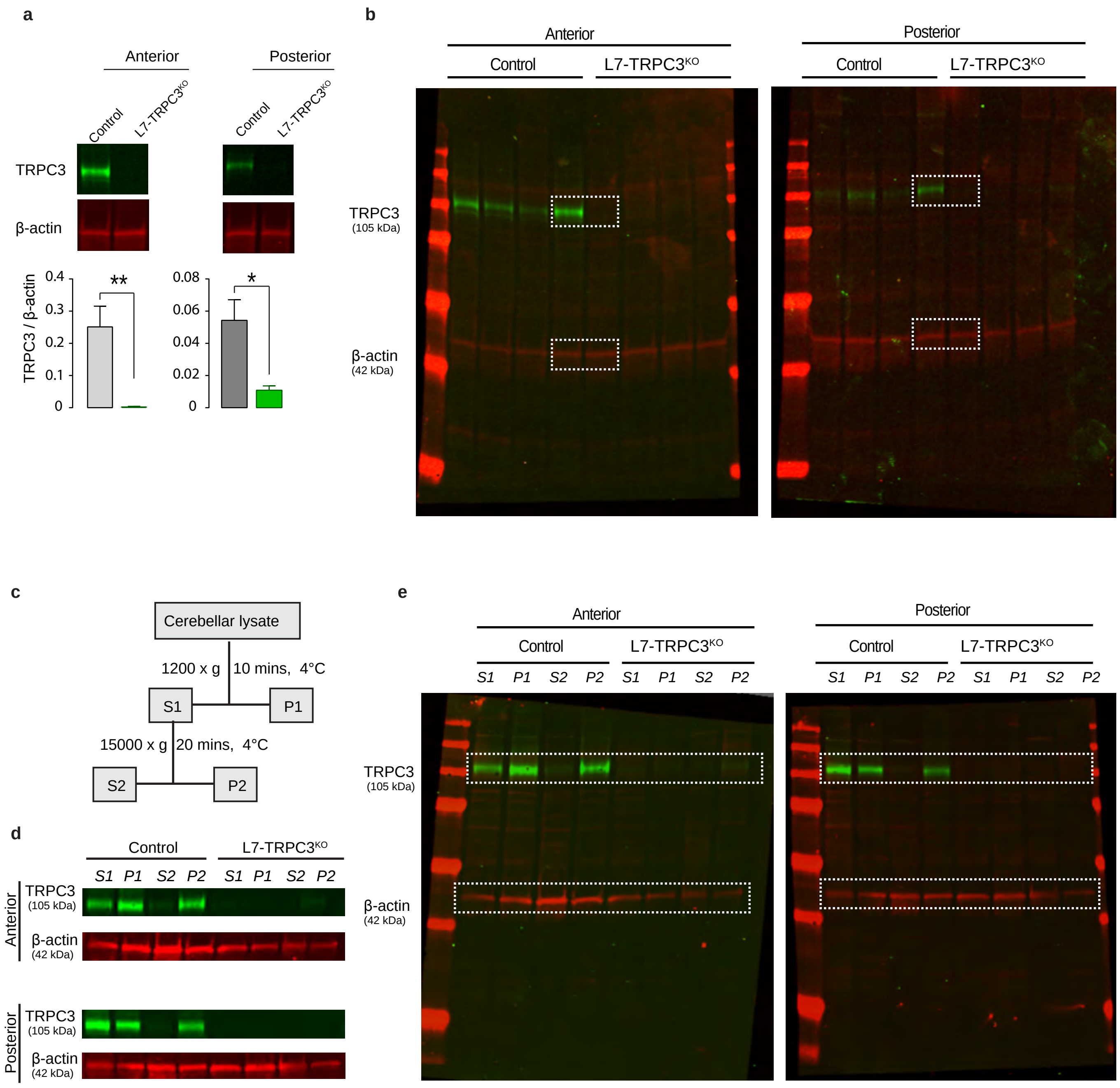


**Supplementary fig.3**

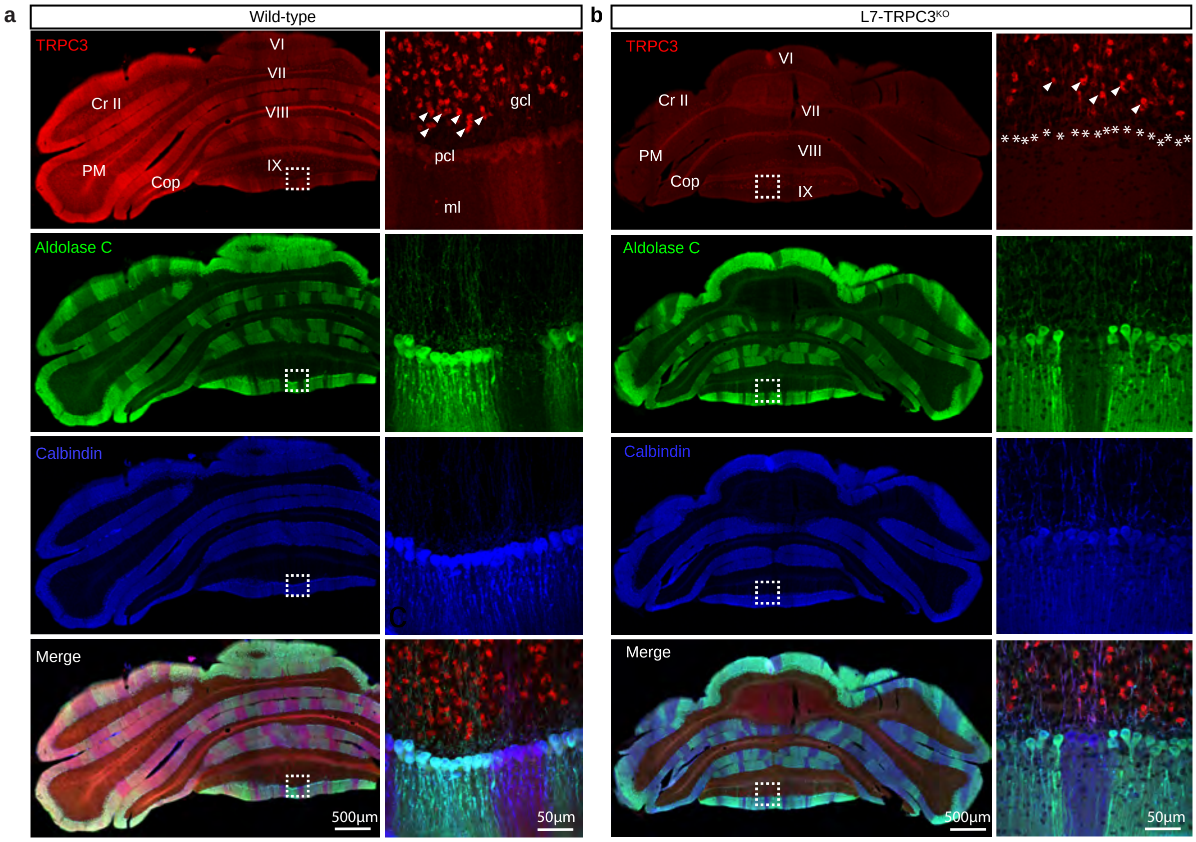


**Supplementary fig.4**

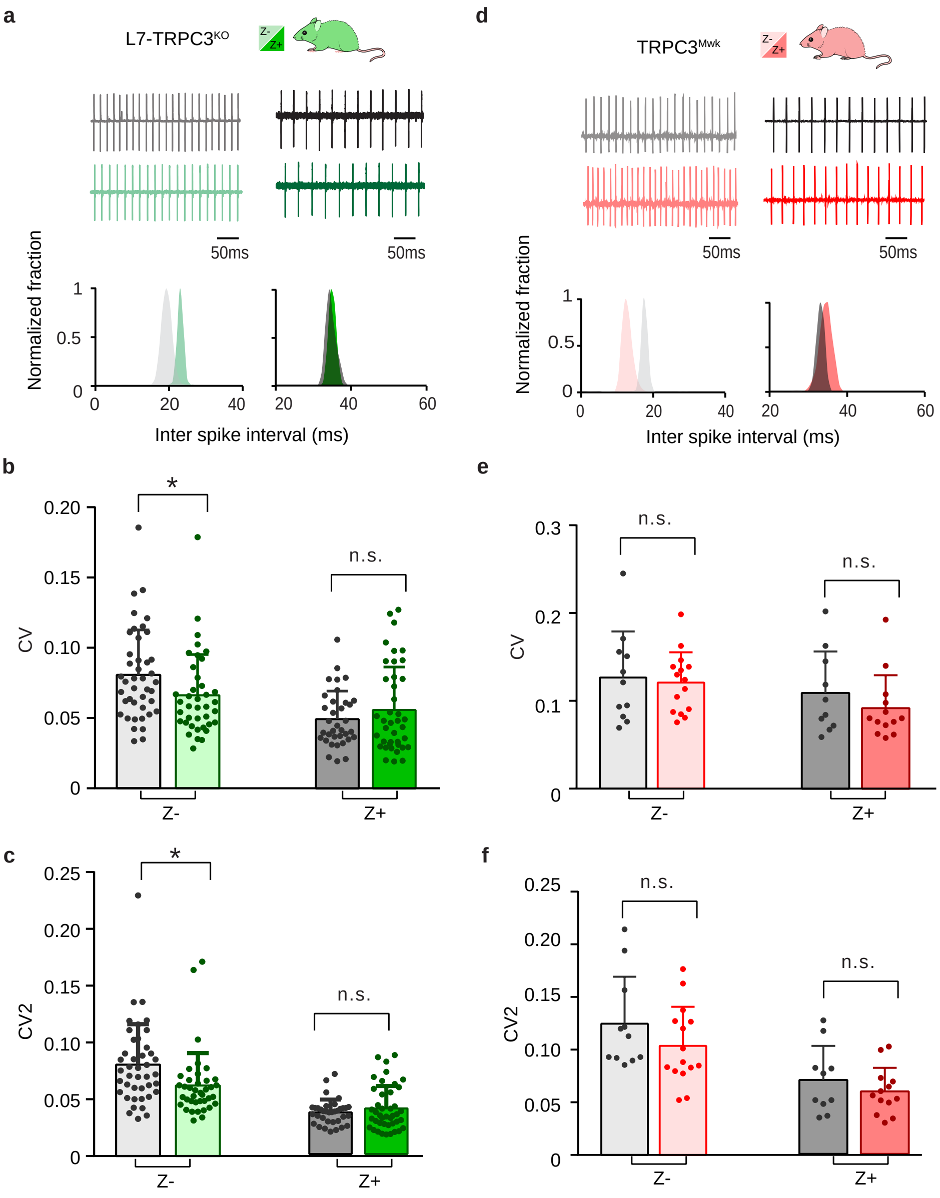


**Supplementary fig.5**

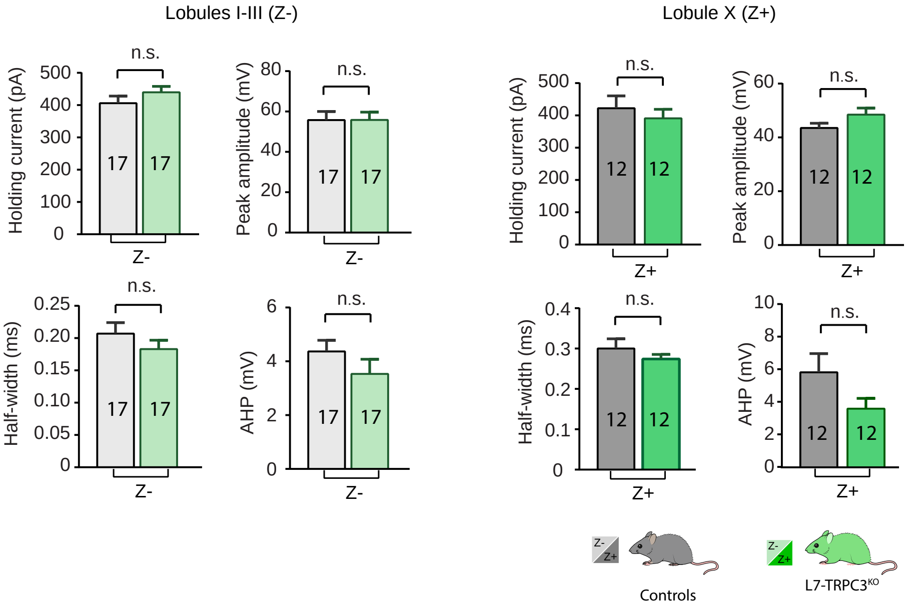


**Supplementary fig.6**

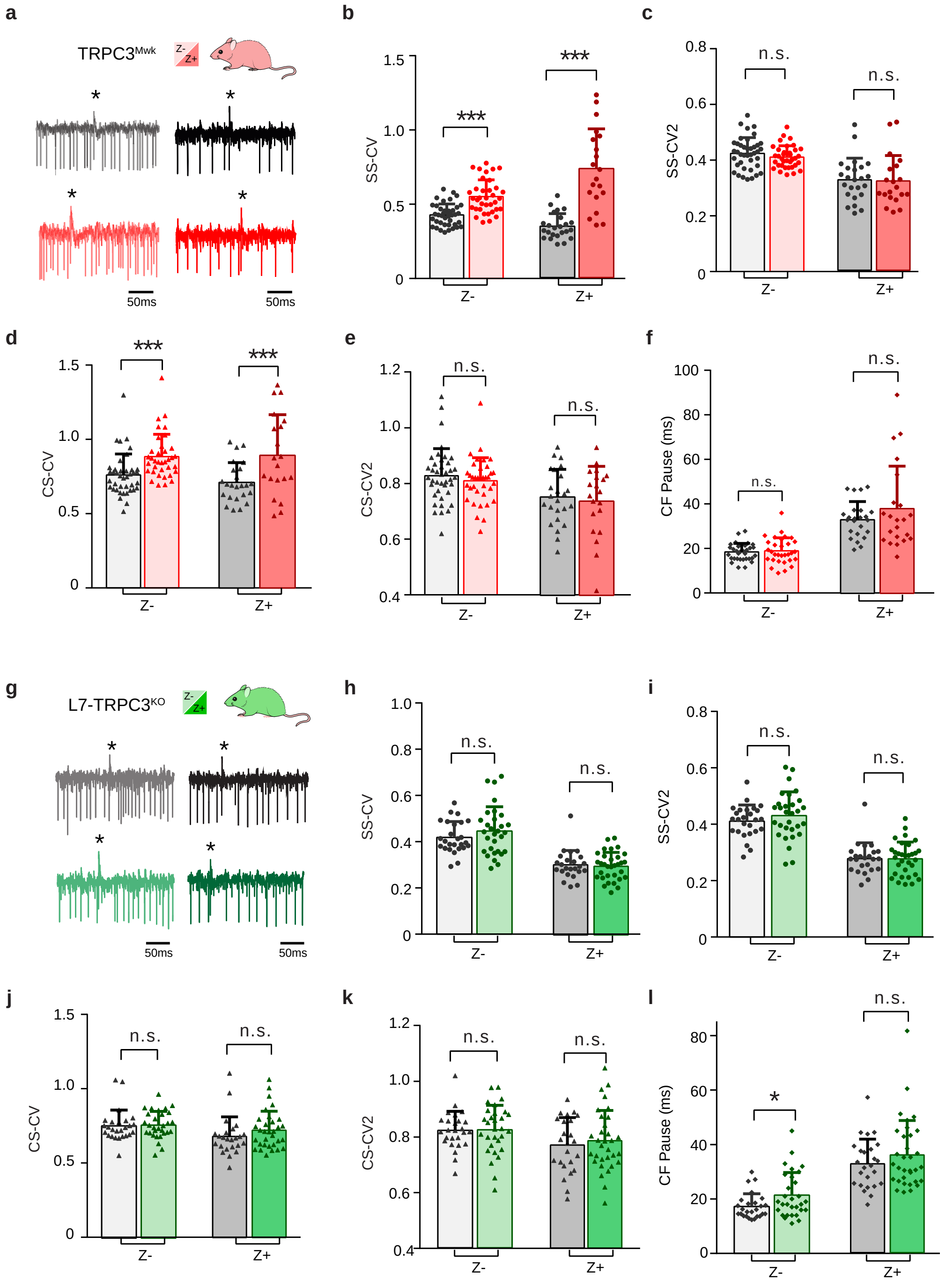


**Supplementary fig.7**

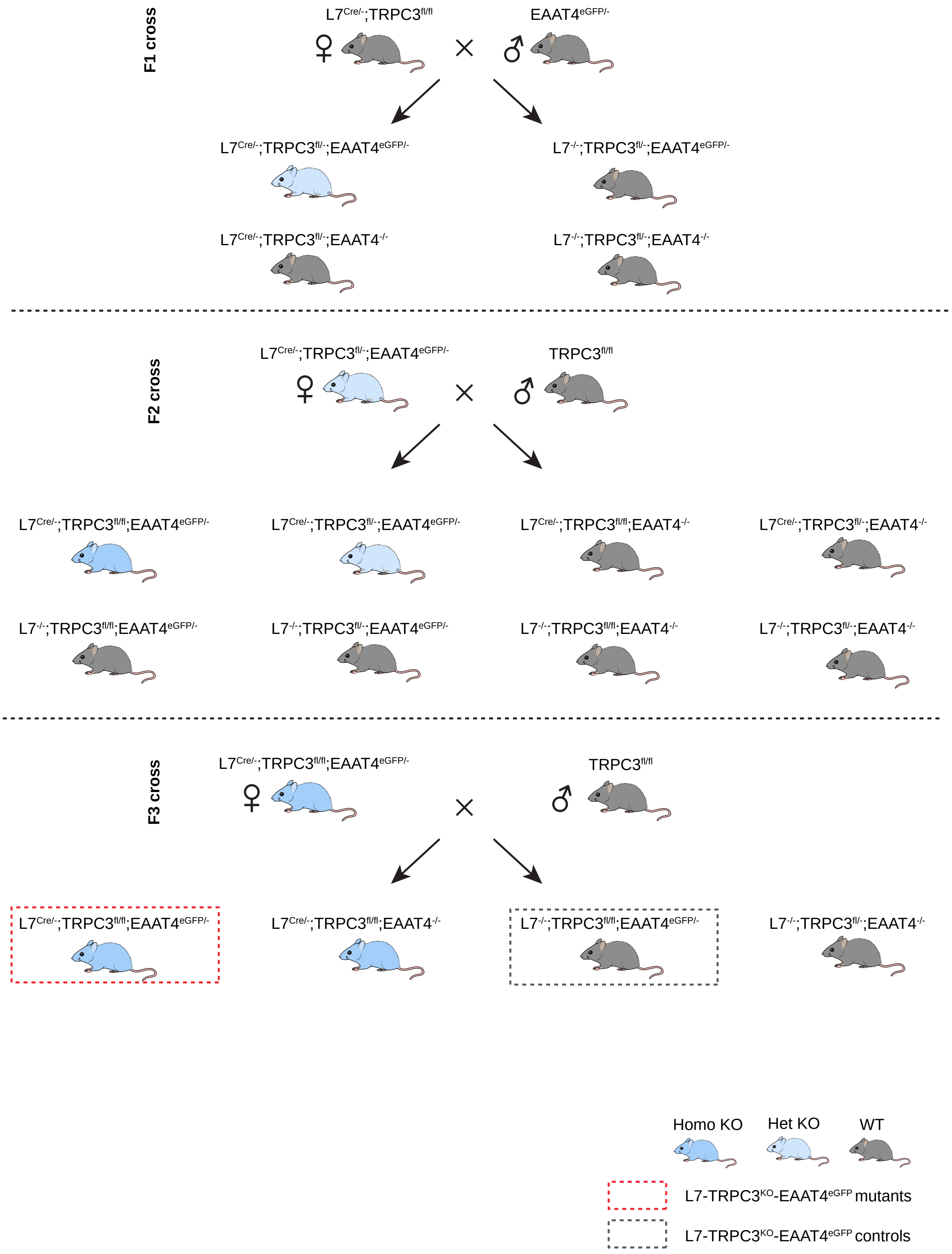


**Supplementary fig.8**

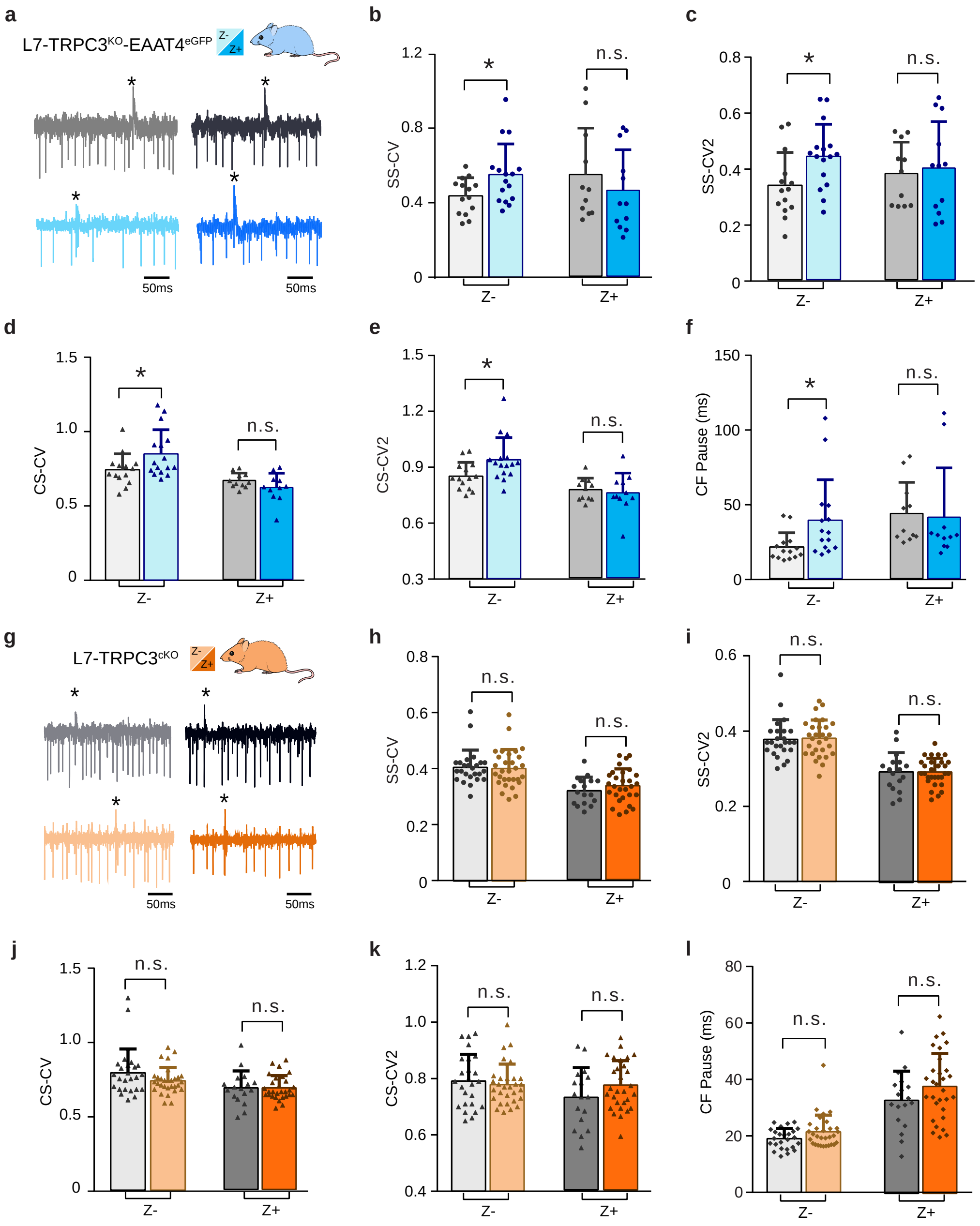


**Supplementary fig.9**

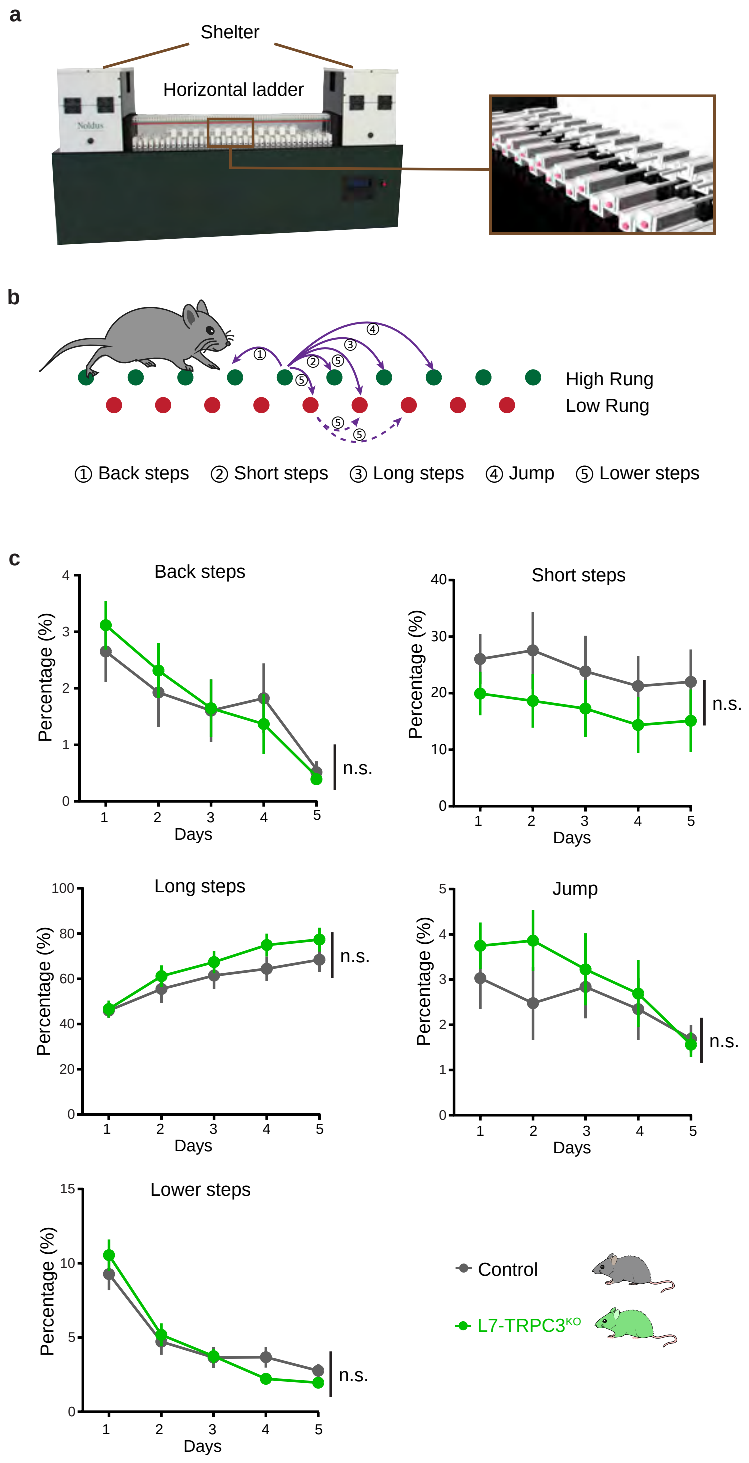


**Supplementary fig.10**

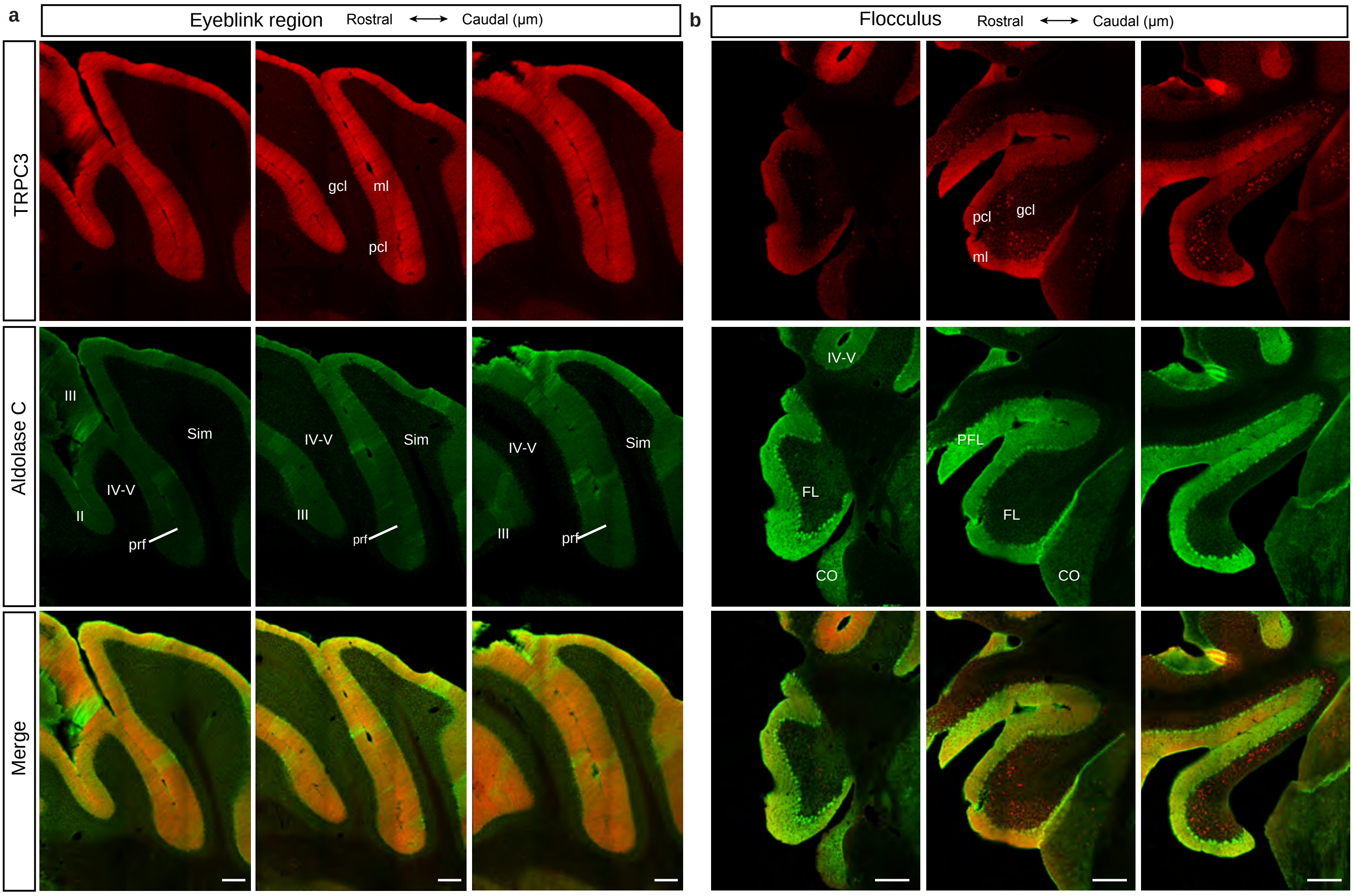


**Supplementary fig.11**

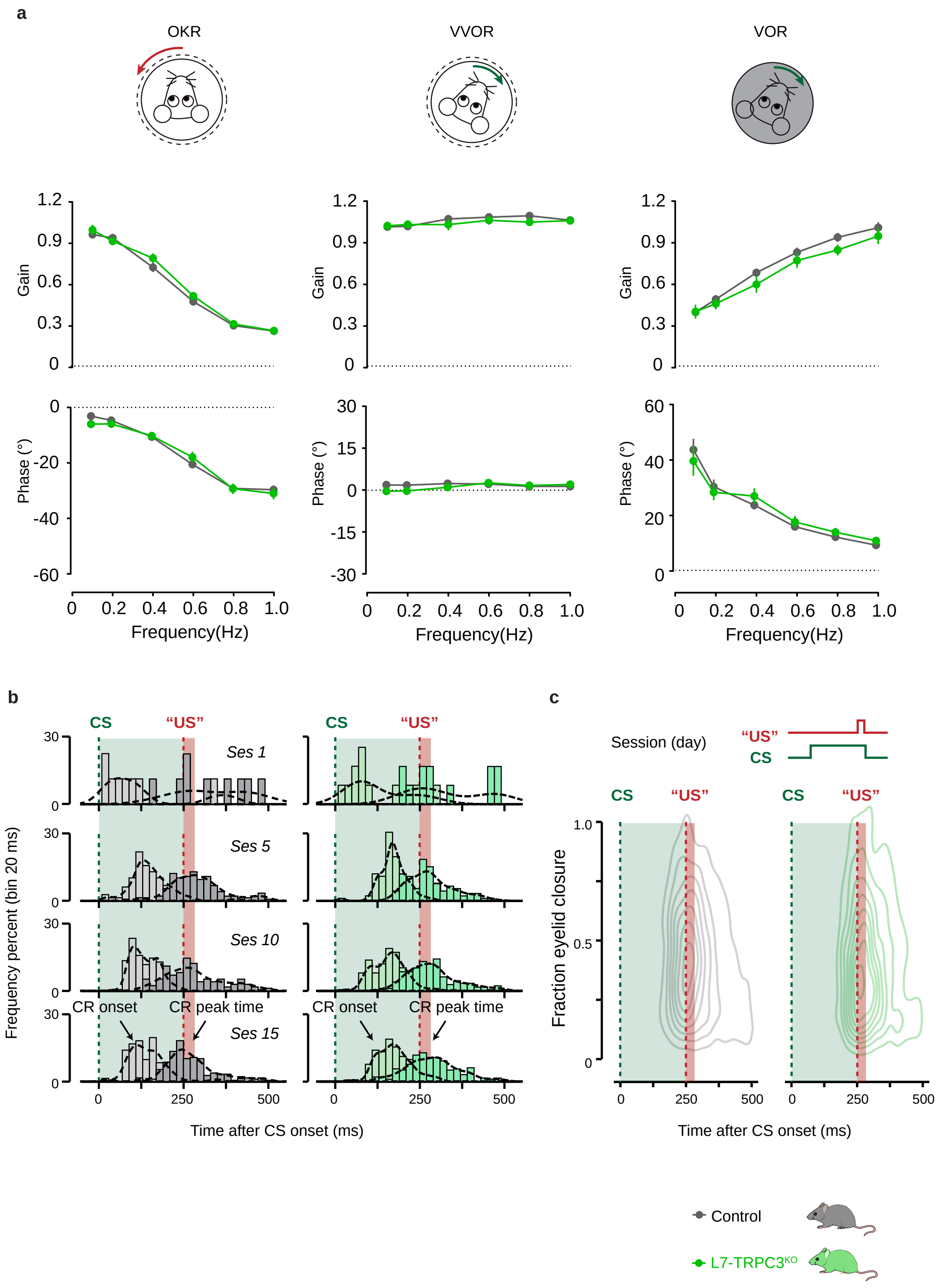


**Supplementary fig.12**

**Supplementary Table 1**

**Summary of the electrophysiological changes**

**Z- PCs**

| Mouse line | In vitro | | | | In vivo | | | | | | |
|---|---|---|---|---|---|---|---|---|---|---|---|
| | *Cell-attach* | | | *Cur-inj* | *Simple spike* | | | *Complex spike* | | | CF-pause |
| | FF | CV | CV2 | FF | FF | CV | CV2 | FF | CV | CV2 | |
| Gain-of-function TRPC3$^{Mwk}$ | ↑ | – | – | N/A | ↑ | ↑ | – | ↓ | ↑ | – | – |
| Loss-of-function L7-TRPC3$^{KO}$ | ↓ | ↓ | ↓ | ↓ | ↓ | – | – | ↓ | – | – | ↑ |
| Loss-of-function L7-TRPC3$^{KO}$-EAAT4$^{eGFP}$ | N/A | | | | ↓ | ↑ | ↑ | ↓ | ↑ | ↑ | ↑ |
| loss-of-function L7-TRPC3$^{cKO}$ | N/A | | | | ↓ | – | – | – | – | – | – |

**Z+ PCs**

| Mouse line | In vitro | | | | In vivo | | | | | | |
|---|---|---|---|---|---|---|---|---|---|---|---|
| | *Cell-attach* | | | *Cur-inj* | *Simple spike* | | | *Complex spike* | | | CF-pause |
| | FF | CV | CV2 | FF | FF | CV | CV2 | FF | CV | CV2 | |
| Gain-of-function TRPC3$^{Mwk}$ | – | – | – | N/A | – | ↑ | – | – | ↑ | – | – |
| Loss-of-function L7-TRPC3$^{KO}$ | – | – | – | – | – | – | – | – | – | – | – |
| Loss-of-function L7-TRPC3$^{KO}$-EAAT4$^{eGFP}$ | N/A | | | | – | – | – | ↓ | – | – | – |
| loss-of-function L7-TRPC3$^{cKO}$ | N/A | | | | – | – | – | – | – | – | – |

**Supplementary Table 2**

***In vitro* acute slice recording kinetics**

**a.** L7-TRPC3$^{KO}$ mice (cell-attached)

| **L7-TRPC3$^{KO}$** | | | | | |
|---|---|---|---|---|---|
| | | *Lobule I-III* | | | |
| | Wild-type | L7-TRPC3$^{KO}$ | t-value | df | P-value(t test) |
| Mice # | 5 | 6 | N/A | N/A | N/A |
| Cells # | 43 | 40 | N/A | N/A | N/A |
| Frequency (Hz) | 55.4±21.8 | 44.1±15.6 | 2.69 | 81 | 0.009 |
| CV | 0.08±0.03 | 0.07±0.03 | 2.14 | 79 | 0.036 |
| CV2 | 0.08±0.04 | 0.06±0.03 | 2.54 | 79 | 0.013 |
| | | *Lobule X* | | | |
| | Wild-type | L7-TRPC3$^{KO}$ | t-value | df | P-value(t test) |
| Mice # | 4 | 10 | N/A | N/A | N/A |
| Cells # | 35 | 36 | N/A | N/A | N/A |
| Frequency (Hz) | 30.8±11.7 | 28.5±9.3 | 0.937 | 64 | 0.352 |
| CV | 0.05±0.02 | 0.06±0.03 | -1.13 | 71 | 0.263 |
| CV2 | 0.04±0.01 | 0.04±0.02 | -0.977 | 67 | 0.332 |

Values are shown as mean±s.d., significant P values are labeled as red.

**b.** TRPC3$^{Mwk}$ mice (cell-attached)

| **TRPC3$^{Mwk}$** | | | | | |
|---|---|---|---|---|---|
| | | *Lobule I-III* | | | |
| | Wild-type | TRPC3$^{Mwk}$ | t-value | df | P-value(t test) |
| Mice # | 2 | 4 | N/A | N/A | N/A |
| Cells # | 11 | 15 | N/A | N/A | N/A |
| Frequency (Hz) | 59.6±14.0 | 84.5±36.2 | -2.43 | 19 | 0.025 |
| CV | 0.13±0.05 | 0.12±0.03 | 0.34 | 24 | 0.735 |
| CV2 | 0.12±0.04 | 0.10±0.04 | 1.32 | 24 | 0.199 |
| | | *Lobule X* | | | |
| | Wild-type | TRPC3$^{Mwk}$ | t-value | df | P-value(t test) |
| Mice # | 2 | 4 | N/A | N/A | N/A |
| Cells # | 10 | 13 | N/A | N/A | N/A |
| Frequency (Hz) | 38.2±16.2 | 36.7±13.0 | 0.242 | 21 | 0.811 |
| CV | 0.11±0.05 | 0.09±0.04 | 0.985 | 21 | 0.336 |
| CV2 | 0.07±0.03 | 0.06±0.02 | 0.960 | 21 | 0.348 |

Values are shown as mean±s.d., significant P values are labeled as red.

**c.** L7-TRPC3$^{KO}$ mice (whole-cell, current injection)

| **L7-TRPC3$^{KO}$** | | | | | |
|---|---|---|---|---|---|
| | | *Lobule I-III* | | | |
| | Widetype | L7-TRPC3$^{KO}$ | t-value | df | P-value(t test) |
| Mice # | 5 | 5 | N/A | N/A | N/A |
| Cells # | 17 | 17 | N/A | N/A | N/A |
| Slope (Hz/100pA) | 19.2±1.1 | 16.0±1.0 | -2.20 | 32 | 0.035 |
| Holding current(pA) | 406±22 | 440±19 | 1.16 | 32 | 0.254 |
| Peak amplitude (mV) | 55.8±4.3 | 55.9±3.9 | 0.02 | 32 | 0.987 |
| Half-width (ms) | 0.21±0.02 | 0.18±0.01 | -1.10 | 32 | 0.279 |
| AHP (mV) | 4.37±0.42 | 3.53±0.55 | -1.20 | 31 | 0.238 |
| | | *Lobule X* | | | |
| | Widetype | L7-TRPC3$^{KO}$ | t-value | df | P-value(t test) |
| Mice # | 4 | 5 | N/A | N/A | N/A |
| Cells # | 12 | 12 | N/A | N/A | N/A |
| Slope (Hz/100pA) | 11.5±1.1 | 10.4±0.5 | -0.95 | 22 | 0.354 |
| Holding current(pA) | 422±38 | 391±28 | -0.66 | 22 | 0.515 |
| Peak amplitude (mV) | 43.5±1.8 | 48.5±2.4 | 1.67 | 22 | 0.109 |
| Half-width (ms) | 0.30±0.02 | 0.27±0.01 | -0.91 | 15 | 0.377 |
| AHP (mV) | 5.81±1.15 | 3.58±0.64 | -1.70 | 22 | 0.108 |

Values are shown as mean±s.e.m., significant P values are labeled as red.

**Supplementary Table 3**

***In vivo* extracellular recording kinetics**

**a.** L7-TRPC3$^{KO}$ mice

| **L7-TRPC3$^{KO}$** | | | | | |
|---|---|---|---|---|---|
| | | **Lobule I-III** | | | |
| | Wild-type | L7-TRPC3$^{KO}$ | t-value | df | P-value(t test) |
| Mice # | 8 | 7 | N/A | N/A | N/A |
| Cells # | 26 | 30 | N/A | N/A | N/A |
| *Simple spike* | | | | | |
| Frequency (Hz) | 88.5±17.4 | 74.4±18.6 | 2.88 | 54 | 0.006 |
| CV | 0.42±0.07 | 0.45±0.10 | -1.17 | 54 | 0.246 |
| CV2 | 0.40±0.06 | 0.42±0.08 | -1.03 | 54 | 0.310 |
| *Complex spike* | | | | | |
| Frequency (Hz) | 1.41±0.35 | 1.16±0.39 | 2.50 | 54 | 0.016 |
| CV | 0.75±0.11 | 0.75±0.09 | -0.217 | 54 | 0.829 |
| CV2 | 0.82±0.07 | 0.82±0.09 | -0.109 | 54 | 0.914 |
| CF-pause(ms) | 17.3±4.7 | 21.4±8.3 | -2.33 | 47 | 0.024 |
| | | **Lobule X** | | | |
| | Wild-type | L7-TRPC3$^{KO}$ | t-value | df | P-value(t test) |
| Mice # | 6 | 8 | N/A | N/A | N/A |
| Cells # | 24 | 32 | N/A | N/A | N/A |
| *Simple spike* | | | | | |
| Frequency (Hz) | 50.0±12.9 | 50.2±15.5 | -0.053 | 54 | 0.958 |
| CV | 0.31±0.06 | 0.30±0.06 | 0.371 | 54 | 0.712 |
| CV2 | 0.28±0.06 | 0.28±0.06 | 0.041 | 54 | 0.967 |
| *Complex spike* | | | | | |
| Frequency (Hz) | 0.98±0.34 | 0.85±0.35 | 1.41 | 54 | 0.164 |
| CV | 0.67±0.13 | 0.71±0.13 | -1.21 | 54 | 0.233 |
| CV2 | 0.77±0.10 | 0.78±0.11 | -0.557 | 54 | 0.580 |
| CF-pause(ms) | 33.5±9.3 | 36.8±13.2 | -1.09 | 56 | 0.281 |

Values are shown as mean±s.d., significant P values are labeled as red.

**b.** TRPC3$^{Mwk}$ mice

| TRPC3$^{Mwk}$ | | | | | |
|---|---|---|---|---|---|
| | | **Lobule I-III** | | | |
| | Wild-type | Trpc3$^{mwk}$ | t-valu | df | p-value(t test) |
| Mice # | 6 | 7 | N/A | N/A | N/A |
| Cells # | 40 | 36 | N/A | N/A | N/A |
| *Simple spike* | | | | | |
| Frequency (Hz) | 89.1±15.3 | 110±22.6 | -4.58 | 60 | $2.41\times10^{-5}$ |
| CV | 0.43±0.07 | 0.55±0.11 | -5.62 | 60 | $5.37\times10^{-7}$ |
| CV2 | 0.42±0.06 | 0.41±0.04 | 1.15 | 74 | 0.254 |
| *Complex spike* | | | | | |
| Frequency (Hz) | 1.16±0.37 | 0.97±0.24 | 2.68 | 68 | 0.009 |
| CV | 0.76±0.14 | 0.88±0.15 | -3.71 | 74 | $4.05\times10^{-4}$ |
| CV2 | 0.83±0.10 | 0.81±0.08 | 0.870 | 74 | 0.387 |
| CF-Pause (ms) | 18.5±3.8 | 19.0±5.8 | -0.420 | 51 | 0.676 |
| | | **Lobule X** | | | |
| | Wild-type | Trpc3$^{mwk}$ | t-value | df | p-value(t test) |
| Mice No. | 5 | 6 | N/A | N/A | N/A |
| Cell No. | 24 | 20 | N/A | N/A | N/A |
| *Simple spike* | | | | | |
| Frequency (Hz) | 45.3±10.4 | 50.6±13.6 | -1.47 | 42 | 0.148 |
| CV | 0.34±0.08 | 0.73±0.26 | -6.22 | 22 | $2.8\times10^{-6}$ |
| CV2 | 0.32±0.08 | 0.31±0.09 | 0.181 | 42 | 0.857 |
| *Complex spike* | | | | | |
| Frequency (Hz) | 0.85±0.27 | 0.71±0.33 | 1.56 | 42 | 0.126 |
| CV | 0.70±0.13 | 0.88±0.27 | -2.71 | 26 | 0.012 |
| CV2 | 0.75±0.10 | 0.73±0.12 | 0.438 | 42 | 0.664 |
| CF-Pause (ms) | 33.1±8.1 | 38.1±19.0 | -1.14 | 28 | 0.263 |

Values are shown as mean±s.d., significant P values are labeled as red.

**c.** L7-TRPC3$^{KO}$-EAAT4$^{eGFP}$ mice

| L7-TRPC3$^{KO}$-EAAT4$^{eGFP}$ | | | | | |
|---|---|---|---|---|---|
| **Z- PCs (Lobule IV- VI)** | | | | | |
| | Wild-type | L7-TRPC3$^{KO}$-EAAT4$^{eGFP}$ | t-value | df | P-value(t test) |
| Mice # | 2 | 3 | N/A | N/A | N/A |
| Cells # | 14 | 16 | N/A | N/A | N/A |
| *Simple spike* | | | | | |
| Frequency (Hz) | 72.7±26.5 | 36.5±23.2 | 3.99 | 28 | $4.4\times10^{-4}$ |
| CV | 0.44±0.10 | 0.55±0.16 | -2.27 | 28 | 0.031 |
| CV2 | 0.34±0.12 | 0.44±0.12 | -2.43 | 28 | 0.022 |
| *Complex spike* | | | | | |
| Frequency (Hz) | 1.18±0.36 | 0.80±0.23 | 3.49 | 28 | 0.002 |
| CV | 0.74±0.11 | 0.85±0.16 | -2.12 | 28 | 0.043 |
| CV2 | 0.85±0.07 | 0.94±0.12 | -2.39 | 28 | 0.024 |
| CF-pause (ms) | 21.1±9.5 | 39.1±27.1 | -2.41 | 18 | 0.027 |
| **Z+ PCs (Lobule IV- VI)** | | | | | |
| | Wild-type | L7-TRPC3$^{KO}$-EAAT4$^{eGFP}$ | t-value | df | P-value(t test) |
| Mice # | 2 | 3 | N/A | N/A | N/A |
| Cells # | 12 | 12 | N/A | N/A | N/A |
| *Simple spike* | | | | | |
| Frequency (Hz) | 33.0±9.8 | 36.6±19.5 | -0.550 | 21 | 0.588 |
| CV | 0.56±0.26 | 0.47±0.22 | 0.873 | 21 | 0.393 |
| CV2 | 0.39±0.11 | 0.41±0.17 | -0.333 | 21 | 0.742 |
| *Complex spike* | | | | | |
| Frequency (Hz) | 1.02±0.20 | 0.74±0.24 | 3.03 | 20 | 0.007 |
| CV | 0.70±0.05 | 0.65±0.10 | 1.44 | 20 | 0.165 |
| CV2 | 0.80±0.06 | 0.78±0.11 | 0.464 | 20 | 0.648 |
| CF-pause (ms) | 46.1±22.0 | 43.4±34.8 | 0.216 | 20 | 0.831 |

Values are shown as mean±s.d., significant P values are labeled as red.

**d.** L7-TRPC3$^{cKO}$ mice

| L7-TRPC3$^{cKO}$ | | | | | |
|---|---|---|---|---|---|
| **Lobule I-III** | | | | | |
| | Wild-type | L7-TRPC3$^{cKO}$ | t-value | df | P-value(t test) |
| Mice # | 4 | 4 | N/A | N/A | N/A |
| Cells # | 25 | 30 | N/A | N/A | N/A |
| *Simple spike* | | | | | |
| Frequency (Hz) | 86.5±10.9 | 72.9±9.1 | 5.05 | 53 | $5.6\times10^{-6}$ |
| CV | 0.40±0.06 | 0.40±0.07 | 0.221 | 53 | 0.826 |
| CV2 | 0.38±0.05 | 0.38±0.05 | -0.222 | 53 | 0.825 |
| *Complex spike* | | | | | |
| Frequency (Hz) | 1.15±0.29 | 1.23±0.31 | -0.940 | 53 | 0.352 |
| CV | 0.78±0.16 | 0.73±0.09 | 1.51 | 53 | 0.136 |
| CV2 | 0.79±0.09 | 0.78±0.07 | 0.570 | 53 | 0.571 |
| CF-pause(ms) | 19.0±3.6 | 21.5±5.9 | -1.83 | 53 | 0.073 |
| **Lobule X** | | | | | |
| | Wild-type | L7-TRPC3$^{cKO}$ | t-value | df | P-value(t test) |
| Mice # | 3 | 4 | N/A | N/A | N/A |
| Cells # | 17 | 29 | N/A | N/A | N/A |
| *Simple spike* | | | | | |
| Frequency (Hz) | 51.6±13.4 | 47.0±11.6 | 1.21 | 44 | 0.234 |
| CV | 0.31±0.05 | 0.33±0.06 | -1.09 | 44 | 0.283 |
| CV2 | 0.29±0.05 | 0.29±0.04 | 0.025 | 44 | 0.980 |
| *Complex spike* | | | | | |
| Frequency (Hz) | 0.98±0.34 | 0.94±0.31 | 0.448 | 44 | 0.656 |
| CV | 0.67±0.11 | 0.68±0.08 | -0.020 | 44 | 0.984 |
| CV2 | 0.73±0.11 | 0.77±0.09 | -1.50 | 44 | 0.140 |
| CF-pause(ms) | 33.0±10.4 | 37.9±11.6 | -1.49 | 46 | 0.143 |

Values are shown as mean±s.d., significant P values are labeled as red.

**Supplementary Table 4 Summary of compensatory eye-movements recording**

**a**, Performance

| Session | OKR | | VOR | | VVOR | |
|---|---|---|---|---|---|---|
| # | WT | MUT | WT | MUT | WT | MUT |
| 1 | 0.96±0.02 | 1.00±0.04 | 1.01±0.02 | 1.02±0.03 | 0.40±0.02 | 0.41±0.05 |
| 2 | 0.94±0.03 | 0.92±0.03 | 1.02±0.02 | 1.03±0.03 | 0.49±0.03 | 0.49±0.04 |
| 3 | 0.73±0.03 | 0.79±0.03 | 1.07±0.01 | 1.03±0.04 | 0.69±0.03 | 0.60±0.06 |
| 4 | 0.48±0.03 | 0.52±0.03 | 1.09±0.02 | 1.06±0.03 | 0.83±0.03 | 0.77±0.05 |
| 5 | 0.30±0.02 | 0.32±0.02 | 1.10±0.02 | 1.05±0.03 | 0.94±0.03 | 0.85±0.04 |
| 6 | 0.26±0.02 | 0.27±0.01 | 1.06±0.02 | 1.06±0.03 | 1.01±0.04 | 0.95±0.06 |

**b**, Gain adaptation

| Session | VOR gain decrease | | VOR gain increase | | OKR gain increase | |
|---|---|---|---|---|---|---|
| # | WT | MUT | WT | MUT | WT | MUT |
| 1 | 0.85±0.03 | 0.78±0.05 | 1.00±0.00 | 1.00±0.00 | 0.94±0.03 | 0.95±0.04 |
| 2 | 0.69±0.03 | 0.62±0.04 | 1.03±0.06 | 1.03±0.09 | 0.94±0.03 | 0.94±0.05 |
| 3 | 0.63±0.05 | 0.58±0.05 | 1.11±0.07 | 1.14±0.10 | 0.93±0.04 | 0.92±0.04 |
| 4 | 0.54±0.06 | 0.52±0.03 | 1.17±0.06 | 1.21±0.07 | 0.90±0.03 | 0.88±0.05 |
| 5 | 0.56±0.03 | 0.54±0.04 | 1.28±0.07 | 1.24±0.09 | 0.87±0.03 | 0.88±0.05 |
| 6 | 0.53±0.03 | 0.48±0.05 | 1.28±0.08 | 1.31±0.09 | 0.78±0.03 | 0.80±0.05 |
| 7 | 0.51±0.04 | 0.49±0.05 | | | | |

**c**, Phase adaptation

| Session | Day1 | | Day2 | | Day3 | | Day4 | | Day5 | |
|---|---|---|---|---|---|---|---|---|---|---|
| # | WT | MUT | WT | MUT | WT | MUT | WT | MUT | WT | MUT |
| 1 | 13±1 | 17±1 | 16±2 | 15±2 | 21±2 | 20±2 | 31±5 | 40±7 | 64±13 | 67±13 |
| 2 | 13±1 | 15±1 | 15±2 | 19±2 | 31±7 | 33±7 | 56±16 | 62±13 | 127±18 | 110±15 |
| 3 | 14±2 | 17±2 | 20±3 | 24±3 | 56±13 | 48±10 | 81±16 | 90±19 | 133±15 | 133±17 |
| 4 | 16±2 | 18±2 | 22±3 | 28±4 | 59±13 | 58±10 | 95±18 | 85±16 | 142±12 | 135±14 |
| 5 | 17±2 | 18±2 | 22±3 | 25±5 | 64±13 | 54±10 | 81±15 | 106±16 | 148±8 | 144±13 |
| 6 | 15±2 | 17±2 | 23±4 | 30±6 | 57±13 | 72±14 | 101±16 | 111±16 | 145±9 | 154±15 |
| 7 | 17±1 | 14±2 | 32±6 | 30±4 | 69±14 | 61±11 | 95±15 | 104±17 | 148±9 | 153±11 |

**d**, Statistics (repeated measures ANOVA )

| Paradigms | Mice # (WT/MUT) | F-value | P-value |
|---|---|---|---|
| OKR (gain / phase) | 12/10 | 0.478 / 0.223 | 0.497 / 0.642 |
| VOR (gain / phase) | 12/10 | 1.13 / 1.24 | 0.300 / 0.279 |
| VVOR (gain / phase) | 12/10 | 0.240 / 0.001 | 0.629 / 0.975 |
| OKR gain increase | 12/10 | 0.010 | 0.922 |
| VOR gain increase | 12/11 | 0.012 | 0.913 |
| VOR gain decrease | 13/11 | 0.252 | 0.621 |
| Phase reversal (Day 5) | 13/11 | 0.006 | 0.942 |

Values are shown as mean±s.e.m., significant P values are labeled as red. MUT is referred to as L7-TRPC3$^{KO}$, and WT is referred to as littermate controls.

**Supplementary Table 5**

**Statistics of the linear mixed-effect model analysis for eyeblink conditioning**

**a**, CR percentage

| **CR percentage** | | | | | |
|---|---|---|---|---|---|
| Session # | Wild-type (N=15) | L7-TRPC3$^{KO}$(N=15) | t-value | df | p-value |
| 1 | 4.82±1.8 | 1.24±0.7 | 0.046 | 28 | 0.963 |
| 2 | 20.8±7.5 | 6.01±1.9 | 1.35 | 28 | 0.189 |
| 3 | 37.9±10.1 | 16.6±7.6 | 2.30 | 28 | 0.029 |
| 4 | 58.9±10.3 | 27.4±8.9 | 3.50 | 28 | 0.002 |
| 5 | 69.0±9.9 | 47.1±10.9 | 2.43 | 28 | 0.022 |
| 6 | 60.0±10.4 | 39.4±8.1 | 2.27 | 28 | 0.031 |
| 7 | 75.4±7.0 | 61.4±8.6 | 1.53 | 28 | 0.138 |
| 8 | 89.2±5.9 | 71.5±8.0 | 1.91 | 28 | 0.066 |
| 9 | 93.5±3.5 | 79.8±7.0 | 1.25 | 28 | 0.220 |
| 10 | 96.0±1.5 | 88.2±6.1 | 0.577 | 28 | 0.569 |
| 11 | 84.5±3.5 | 78.2±6.2 | 0.701 | 28 | 0.489 |
| 12 | 88.1±4.7 | 89.2±4.3 | -0.080 | 28 | 0.937 |
| 13 | 96.5±1.7 | 90.6±2.5 | 0.667 | 28 | 0.510 |
| 14 | 93.0±2.2 | 91.5±3.5 | 0.150 | 28 | 0.882 |
| 15 | 88.1±7.5 | 82.2±8.0 | 0.295 | 28 | 0.770 |

**b**, Fraction eyelid closure

| **Fraction eyelid closure** | | | | | |
|---|---|---|---|---|---|
| Session # | Wild-type(N=15) | L7-TRPC3$^{KO}$(N=15) | t-value | df | p-value |
| 1 | 0.016±0.005 | 0.012±0.002 | -0.404 | 28 | 0.689 |
| 2 | 0.071±0.027 | 0.015±0.004 | 0.387 | 28 | 0.702 |
| 3 | 0.184±0.055 | 0.057±0.026 | 1.75 | 28 | 0.092 |
| 4 | 0.297±0.064 | 0.118±0.049 | 2.46 | 28 | 0.020 |
| 5 | 0.461±0.108 | 0.213±0.059 | 3.64 | 28 | 0.001 |
| 6 | 0.279±0.06 | 0.194±0.051 | 1.25 | 28 | 0.223 |
| 7 | 0.362±0.056 | 0.314±0.077 | 0.508 | 28 | 0.615 |
| 8 | 0.552±0.074 | 0.425±0.081 | 1.52 | 28 | 0.139 |
| 9 | 0.614±0.082 | 0.401±0.056 | 2.75 | 28 | 0.010 |
| 10 | 0.658±0.071 | 0.533±0.078 | 1.38 | 28 | 0.179 |
| 11 | 0.543±0.078 | 0.462±0.075 | 1.51 | 28 | 0.141 |
| 12 | 0.541±0.081 | 0.513±0.06 | 0.907 | 28 | 0.372 |
| 13 | 0.686±0.066 | 0.605±0.063 | 1.38 | 28 | 0.178 |
| 14 | 0.577±0.055 | 0.535±0.063 | 0.353 | 28 | 0.727 |
| 15 | 0.700±0.091 | 0.518±0.084 | 1.47 | 28 | 0.153 |

Values are shown as mean±s.e.m., P values were all FDR corrected for multiple comparisons, significant P values are labeled as red.

**Supplementary Table 6 Summary of Erasmus Ladder performance**

**a**, Performance

| Session # | Back steps % | | Short steps % | | Long steps % | |
|---|---|---|---|---|---|---|
| | WT | MUT | WT | MUT | WT | MUT |
| 1 | 3.24±0.43 | 2.65±0.54 | 19.9±3.9 | 26.1±4.4 | 46.6±3.8 | 45.9±3.4 |
| 2 | 2.42±0.48 | 1.93±0.61 | 18.6±4.7 | 27.6±6.8 | 61.2±4.8 | 55.5±6.1 |
| 3 | 1.76±0.51 | 1.60±0.55 | 17.3±5.0 | 23.8±6.3 | 67.4±5.0 | 61.5±6.0 |
| 4 | 1.45±0.53 | 1.82±0.62 | 14.4±4.9 | 21.3±5.3 | 74.9±5.0 | 64.4±5.5 |
| 5 | 0.47±0.08 | 0.57±0.14 | 15.1±5.5 | 22.0±5.7 | 77.3±5.3 | 68.4±5.4 |

| Session # | Jump % | | Lower steps % | |
|---|---|---|---|---|
| | WT | MUT | WT | MUT |
| 1 | 3.75±0.51 | 3.03±0.68 | 10.5±1.1 | 9.3±1.1 |
| 2 | 3.86±0.67 | 2.48±0.81 | 5.2±0.8 | 4.7±0.9 |
| 3 | 3.23±0.80 | 2.84±0.70 | 3.8±0.5 | 3.7±0.7 |
| 4 | 2.69±0.74 | 2.34±0.68 | 2.2±0.3 | 3.7±0.7 |
| 5 | 1.56±0.28 | 1.69±0.30 | 2.0±0.2 | 2.8±0.5 |

**b**, Statistics (repeated measures ANOVA)

| Step types | Mice # (WT/MUT) | F-value | p-value |
|---|---|---|---|
| Back steps | 16/16 | 0.011 | 0.917 |
| Short steps | 16/16 | 0.980 | 0.330 |
| Long steps | 16/16 | 0.923 | 0.344 |
| Jump | 16/16 | 0.464 | 0.501 |
| Lower step | 16/16 | 0.012 | 0.913 |

Values are shown as mean±s.e.m., MUT is referred to as L7-TRPC3$^{KO}$, and WT is referred to as littermate controls.
